# CXCL10^+^ peripheral activation niches couple preferred sites of Th1 entry with optimal APC encounter

**DOI:** 10.1101/2020.10.04.324525

**Authors:** Hen Prizant, Nilesh Patil, Seble Negatu, Alexander McGurk, Scott A. Leddon, Angela Hughson, Tristan D. McRae, Yu-Rong Gao, Alexandra M Livingstone, Joanna R Groom, Andrew D Luster, Deborah J Fowell

## Abstract

Correct positioning of T cells within infected tissues is critical for T cell activation and pathogen control. Upon tissue entry, effector T cells must efficiently locate antigen presenting cells (APC) for peripheral activation. We reveal that tissue entry and initial peripheral activation of Th1 effector T cells are tightly linked to perivascular positioning of chemokine-expressing APCs. Dermal inflammation induced tissue-wide *de novo* generation of discrete perivascular CXCL10^+^ cell clusters, enriched for CD11c^+^MHC-II^+^ monocyte-derived dendritic cells. These chemokine clusters were ‘hot spots’ for both Th1 extravasation and activation in the inflamed skin. CXCR3-dependent Th1 localization to the cluster micro-environment prolonged T-APC interactions and boosted function. Both the frequency and range of these clusters were enhanced via a Th1-intrinsic, IFNγ-dependent positive feedback loop. Thus, the perivascular CXCL10^+^ clusters act as initial peripheral activation niches, optimizing controlled activation broadly throughout the tissue by coupling Th1 tissue entry with enhanced opportunities for Th1-APC encounter.

## INTRODUCTION

The spatiotemporal control of effector T cell activation at sites of inflammation and infection is critical for precise delivery of effector molecules while limiting collateral damage. For pathogen control within infected tissues, CD4^+^ effector T cells must locate cognate antigen-bearing antigen presenting cells (APCs) for peripheral re-activation of effector functions (Mandl et al., 2014; Masopust and Schenkel, 2013). The frequency of relevant antigen-bearing APCs within an infected site appears to be limiting (Egen et al., 2011) therefore the range, and efficiency, of tissue search for ligand by CD4^+^ T cell effectors will play an important role in successful anti-pathogen immunity.

Models for how T cells efficiently search the inflamed tissue have been informed by intravital imaging studies that have revealed a surprising degree of apparent random movement of T cells at infection and tumor sites (Filipe-Santos et al., 2009; Harris et al., 2012; Mrass et al., 2006; Overstreet et al., 2013), guided on a micro-anatomical scale by local chemical and physical cues (Gaylo et al., 2016b; Krummel et al., 2016; Sarris and Sixt, 2015) and a paucity of ligand-bearing targets, APC (Egen et al., 2011). The scope of T cell search will be likely influenced by the point of tissue entry, the distribution pattern of relevant APC and the spatial relationship between antigen presentation and the pathogen itself. However, the efficiency of immune responses can be enhanced by reducing random distribution through clustering targets in regions of higher density. In inflamed peripheral tissues, small self-organizing clusters of T cells with innate effectors have been described: perivascular clustering of APCs and CD8^+^ T cells in the skin (Natsuaki et al., 2014), clustering of APCs and Th2 cells in the asthmatic lung (Veres et al., 2017) and macrophage clusters that sustain resident tissue memory T cells (Iijima and Iwasaki, 2014). The positional cues that shape the recruitment, composition and retention of T cells within these niches are not fully understood.

Chemokines, and other chemoattractants, play important roles in the spatial organization of lymphoid tissues and in the movement of leukocytes within and between lymphoid and peripheral tissues (Griffith et al., 2014; Krummel et al., 2016). How chemokines direct the accumulation of T cells to particular microanatomical regions within tissues remains elusive (Sarris and Sixt, 2015). Chemokine sources within inflamed tissues are speculated to locally tune T cell migration in a form of “guided randomness” that may result in iterative small directional changes that result in location-specific T cell accumulation (Weber et al., 2013). This may be achieved by precise spatial display of chemokines through cell- or ECM-associated heparin sulfate immobilization (Schumann et al., 2010; Sorokin, 2010). Additional chemokine control of positioning comes from the dynamic regulation of chemokine receptor sensitivity that can lead to chemokine-mediated arrest at high chemokine concentrations (Lammermann and Kastenmuller, 2019; Sarris et al., 2012). Thus, chemokines may serve as both ‘stop’ and ‘go’ signals within tissues. For example, acute blockade of the chemokines CXCL9 and CXCL10 has been shown to halt T cell interstitial migration (Gaylo-Moynihan et al., 2019; Harris et al., 2012), while T cells deficient in the CXCL9/10-receptor CXCR3 have defects in ‘stopping’ at viral infection foci (Hickman et al., 2015). How T cells integrate chemokine signals with other positional cues such as cognate antigen is unclear.

In this report we have used intravital multiphoton microscopy (IV-MPM) and a CXCL9/CXCL10 reporter system (Groom et al., 2012) to define the spatial and temporal relationship between CXCR3 chemokine availability and the dynamic range of CD4^+^ Th1 cell migration *in vivo*. We reveal that the first step in peripheral activation of Th1 effector T cells is tightly linked to tissue entry through perivascular positioning of chemokine-expressing APCs. Following immunization or infection of the skin, CXCL9 and CXCL10 expression was microanatomically restricted to distinct perivascular clusters of APCs, enriched for moDCs, that ‘marked’ a previously unrecognized spatial preference for Th1 cell accumulation. These clusters were distributed throughout the inflamed ear pinna, often 1,000’s of microns from the immunization site, forming distinct CXCL<u>10</u>^+^ <u>p</u>eripheral <u>ac</u>tivation (PAC-10) niches that facilitate tissue-wide immune activation. The CXCL10^+^ chemokine-rich clusters served to nucleate the recruitment of CXCR3^+^ Th1 cells from the bloodstream and optimize durable Th1 encounters with cognate-Ag bearing APC for initial peripheral activation. Th1 dynamics in the PAC-10 niches were regulated by CXCR3-dependent localization to the chemokine-rich cluster, and by antigen-dependent retention within the cluster. Moreover, we find a key role for IFNγ-producing CD4^+^ T cells themselves in shaping the composition, number and position of the PAC-10 niches. These findings suggest incoming effector T cells play a crucial role in dictating the magnitude and tissue-range of initial activation at inflamed sites through martialing chemokine-rich peripheral activation niches.

## RESULTS

### Perivascular CXCL10-expressing cell clusters define sites of spatial preference for Th1 cell accumulation within the inflamed skin

The CXCR3 chemokines CXCL9 and CXCL10 are strongly upregulated in response to many immune challenges. In response to immunization with protein antigens and Complete Freund’s Adjuvant (CFA) CXCL10 is one of the dominant chemokines expressed at the site of cutaneous inflammation (Fig 1a), however the phenotype and spatial distribution of CXCL10-expressing cells in this context was not known. To identify the location and phenotype of cells expressing CXCR3 chemokines within the inflamed skin we utilized the REX3 reporter mouse(Groom et al., 2012); in which transcriptional expression of CXCL9 and CXCL10 are linked to RFP and BFP respectively. REX3 mice were intradermally (i.d.) immunized with OVA protein emulsified in CFA (OVA/CFA) in the ear pinna and the kinetics of CXCL9 and CXCL10 expression determined on whole ear cell suspensions by flow cytometric analysis. BFP-CXCL10 was rapidly and strongly induced within the hematopoietic compartment (Fig. 1b; S1A, B), with fewer cells expressing RFP-CXCL9 (Fig. 1b). The vast majority of RFP-CXCL9^+^ cells co-expressed BFP-CXCL10, with very few single positive RFP-CXCL9^+^ cells (Fig. S1A, B). We used non-invasive intra-vital multi photon microscopy (IV-MPM) to visualize the precise location of the BFP-CXCL10^+^ cells in 3D within the inflamed skin. BFP and RFP expression was found in distinct perivascular regions of the inflamed tissue. *In-vivo* labeling of blood vessels revealed that the BFP-CXCL10^+^ cell clusters appeared in close proximity to CD31^+^ vessels (Fig. 1c, d). Computational analysis of the distance of each BFP-CXCL10^+^ cell to the nearest CD31^+^ vessel in 3D found that most BFP-CXCL10^+^ cells were either in contact or close (within 10μm) to a CD31^+^ vessel (Fig. 1e-g, Video S1). The average distance of BFP-CXCL10^+^ cells from the CD31^+^ vessels was significantly shorter than the average distance of cells if they were to be distributed randomly within the same field (Fig. 1f). Similar proximity analysis of BFP-CXCL10^+^ cells to LYVE-1^+^ lymphatic vessels did not reveal an association of BFP-CXCL10^+^ cells with lymphatics (Fig. 1g, Fig. S1C, D). The co-expressing RFP-CXCL9^+^ and BFP-CXCL10^+^ cells were also found to be perivascular. Thus, following immunization, clusters of CXCL9 and CXCL10 expressing cells are spatially restricted to perivascular areas.

**Figure 1.**
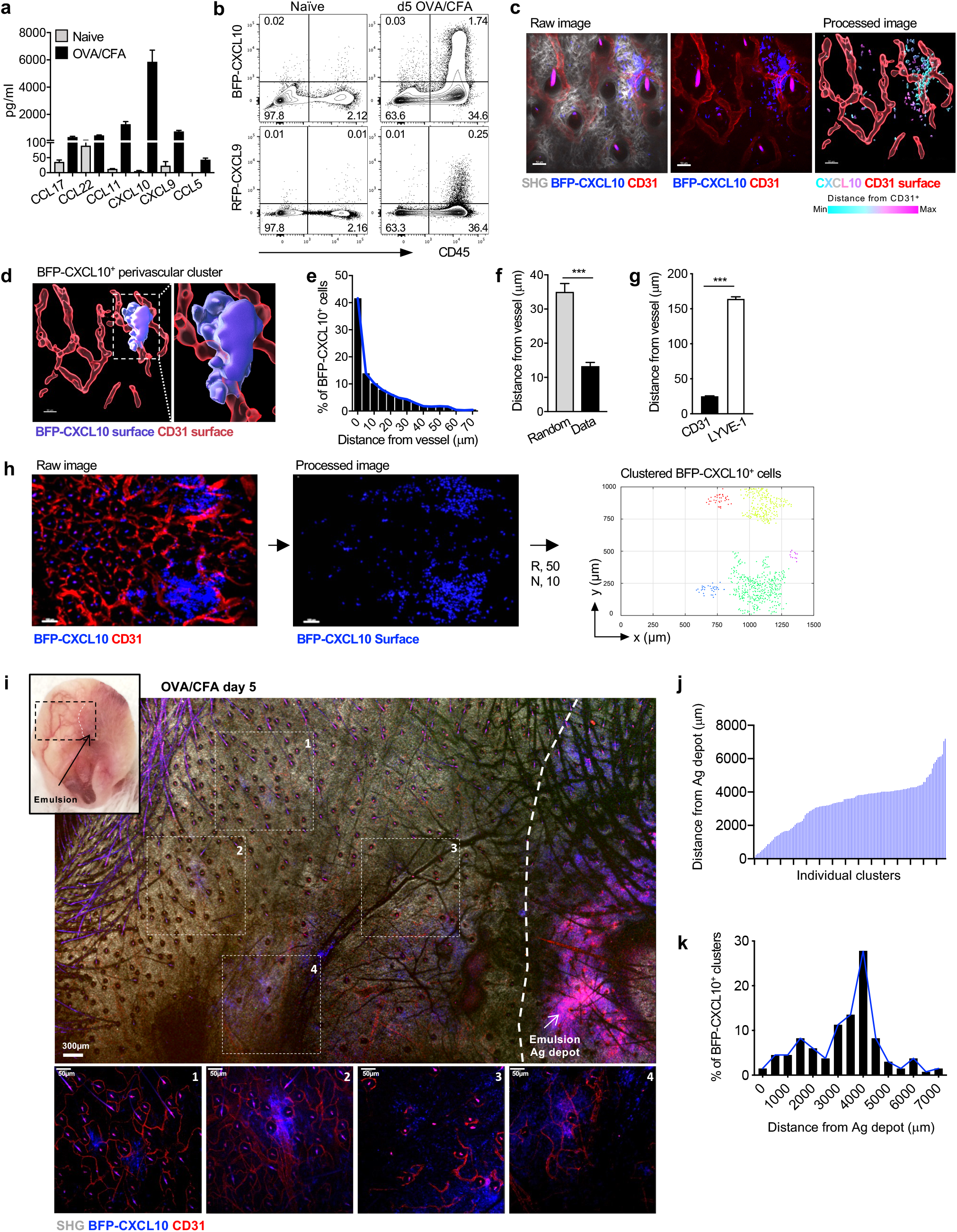
Th1 Accumulation at perivascular BFP-CXCL10^+^ clusters within the inflamed ear. **(a)** Chemokine protein levels from d3 OVA/CFA immunized WT ears. **(b-j)** d5 OVA/CFA dermal immunization of REX3 mice. **(b)** Frequency of BFP-CXCL10 and RFP-CXCL9 expressing hematopoietic cells. Representative from 5 independent experiments, >20 REX mice per group. **(c)** Distribution of BFP-CXCL10^+^ cells (blue) and CD31^+^ vessels (red) by IV-MPM in the dermis: gray, SHG; scale bar 50µm. Processed image: CD31^+^ vessels (red) reconstructed and volumetrically rendered in 3D using Imaris, BFP-CXCL10^+^ cells color coded by distance from vessel. Representative maximal z-projection image, 4 independent experiments, 12 ears, >25 imaging volumes. **(d)** 3D surface rendering of BFP-CXCL10^+^ perivascular cluster (Imaris). **(e)** Frequency distribution of BFP-CXCL10^+^ cells from CD31^+^ vessels. **(f)** BFP-CXCL10^+^ cell distance from CD31^+^ vessels (Data) compared to random distribution in same imaging volume (Random). **(g)** BFP-CXCL10^+^ cell distance from CD31^+^ and LYVE-1^+^ vessels. **(h)** Tiled image of 6 imaging fields of 512 (x) x 512 (y) x 60 (z) μm, scale bar 100µm, raw and processed images as in (c). Right panel, semi-automated 3D cluster reconstruction using DBSCAN-based algorithm: R, radius from cell centroid (μm); N, number of neighbors within the radius. Each dot represents a single detected BFP-CXCL10^+^ cell within identified clusters, with each cell in the same cluster designated a cluster-specific color. **(i)** Tissue-wide cluster distribution relative to the OVA/CFA emulsion (Ag-depot), IV-MPM tiled image of region highlighted in upper left picture using a 10x objective, scale bar 300 μm. Individual images (1-4) acquired with 25x objective (below), scale bar 50 μm. **(j)** Individual cluster distance and **(k)** frequency of BFP-CXCL10^+^ cell distance from Ag depot. Cluster identified using DBSCAN-based algorithm as in (h) and distances calculated between the center coordinates of the identified cluster and emulsion using Pythagorean theorem. 3 independent experiments, 133 clusters, >20 imaging volumes.

For unbiased analysis of T cell migration and positioning relative to these chemokine clusters we employed semi-automated density-based spatial clustering of application with noise (DBSCAN)(Ester et al., 1996) (Fig. S2). This computational analysis enabled the accurate 3D identification and reconstruction of high density BFP-CXCL10^+^ cell clusters in the inflamed skin (Fig. 1h, Fig. S2). The BFP-CXCL10^+^ clusters were not observed in the naïve ear dermis prior to immunization (Fig. S2). To also determine whether perivascular clustering of BFP-CXCL10^+^ cells was observed following cutaneous infection, we analyzed BFP-CXCL10 distribution in the *Leishmania major* (*L. major*)-infected dermis. *L. major* infection leads to the induction of a number of chemokines including CXCL9 and CXCL10 and these chemokine producing cells were also found distributed perivascularly (Fig. S3).

The use of OVA/CFA immunization enabled us to determine the position of the CXCL10^+^ perivascular clusters relative to the antigen depot. We were able to identify the OVA/CFA emulsion site (Ag depot) and examine a large area of the inflamed ear pinna by MPM (Fig. 1i). Unexpectedly, BFP-CXCL10^+^ cells were found in distinct clusters scattered throughout the tissue, often 1,000’s of μm from the emulsion site (Fig. 1i). Analysis of BFP^+^ clusters using our unbiased clustering tool, revealed a broad distribution of clusters across the tissue (Fig. 1j) with the majority of clusters some 4,000 μm from the antigen depot (Fig. 1k). Thus, immunization in the skin induces the de-novo formation of distinct perivascular clusters of CXCR3-chemokine producing cells with far-reaching distribution.

### Th1 cells enter the skin at sites of BFP-CXCL10^+^ clusters and accumulate in a CXCR3-mediated manner

Once clusters were defined, we were able to quantitate the positioning of Th1 cells relative to the chemokine cluster. *In vitro*-generated, adoptively transferred Th1 cells preferentially accumulated at sites of CXCL10^+^ cell perivascular clusters (Fig 2a, b). To assess whether chemokines CXCL9 or CXCL10 were playing an active role in Th1 accumulation, WT and CXCR3 deficient (CXCR3 KO) fluorescently-labelled Th1 cells were co-transferred and their position was assessed relative to the BFP-CXCL10^+^ perivascular clusters. CXCR3 KO Th1 cells failed to accumulate within BFP-CXCL10^+^ clusters (Fig. 2 c, d; Video S2). Similarly, OT-II Th2 cells, that naturally fail to upregulate the expression of CXCR3(Griffith et al., 2014; Sallusto and Lanzavecchia, 2000), were also not restricted to the BFP-CXCL10^+^ clusters in the OVA/CFA dermis (Fig. S4). Notably, CXCR3-dependent Th1 accumulation within BFP-CXCL10^+^ clusters did not occur in the absence of cognate peptide: OVA-specific Th1 cells did not accumulate within the BFP-CXCL10^+^ clusters formed in the KLH/CFA immunized dermis (Fig 2e, f; Fig. S5, Video S3), suggesting chemokine signals alone are not sufficient to position T cells in the perivascular clusters. Thus, in the inflamed skin we observe the *de novo* induction of perivascular BFP-CXCL10^+^ cell clusters of hematopoietic origin that marked an unrecognized spatial preference for antigen-dependent CXCR3^+^ Th1 cell accumulation.

**Figure 2.**
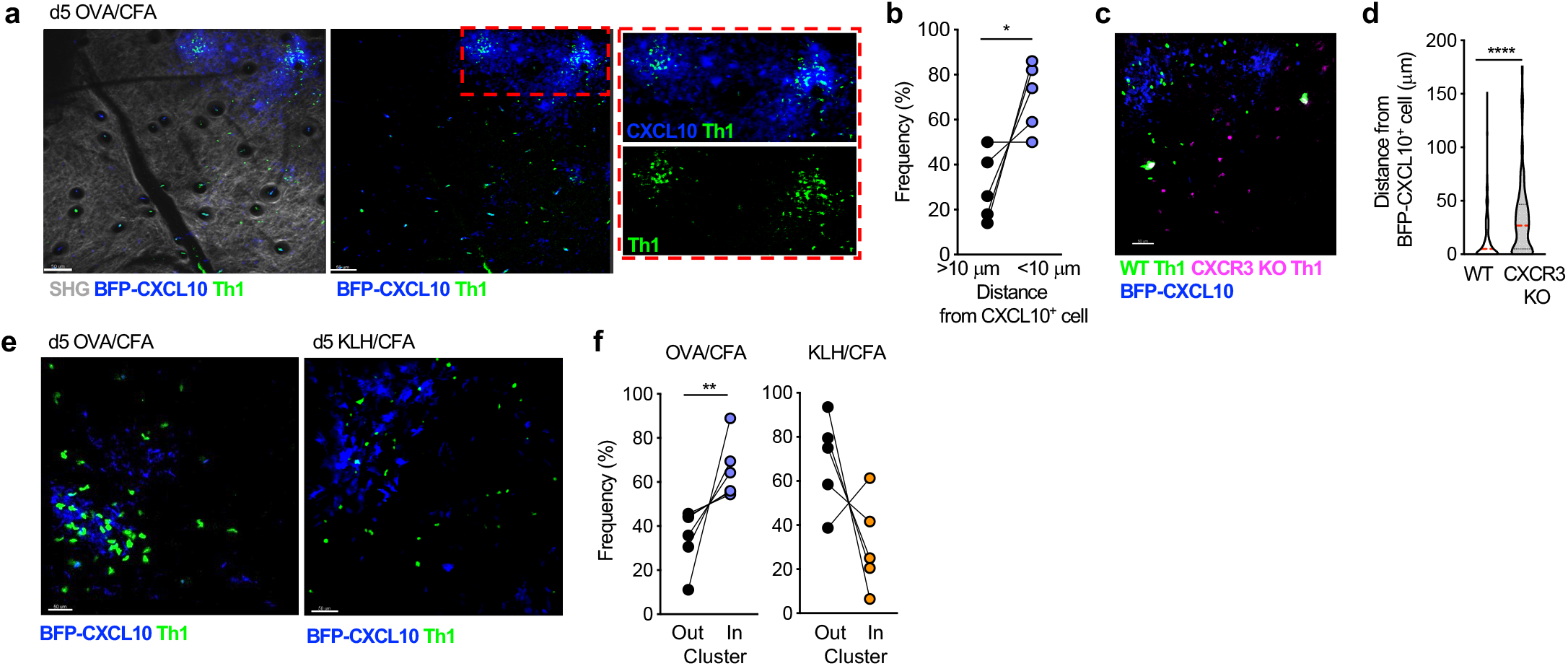
Th1 cell spatial preference. OT-II Th1 cells were transferred i.v. into OVA/CFA immunized REX3 mice and their position relative to the BFP-CXCL10^+^ clusters determine on d5 of immunization by IV-MPM of the inflamed dermis. **(a)** Adoptively transferred OT-II Th1 cells (green) localized to BFP-CXCL10^+^ clusters (blue). Maximal z-projection by IV-MPM: gray, SHG; scale bar 50µm. **(b)** Frequency of OT-II Th1 cells within 10μm of BFP-CXCL10^+^ cells, paired data points from the same imaging field. **(c, d)** Adoptively co-transferred WT (green) and CXCR3 KO (magenta) OT-II Th1 cell position relative to BFP-CXCL10^+^ cells (blue). Representative maximal z-projection by IV-MPM, scale bar 50µm. **(d)** WT and CXCR3-KO OT-II Th1 cell distance from BFP-CXCL10^+^ cells. Pooled data from 3 independent experiments, >15 imaging volumes. **(e, f)** Immunization with cognate (OVA/CFA) or non-cognate (KLH/CFA) antigen and transfer of OVA-specific OT-II Th1 cells. **(e)** Representative maximal z-projection by IV-MPM, scale bar 50µm and **(f)** frequency of OT-II Th1s (green) relative to BFP-CXCL10^+^ clusters (blue). Data from 3 independent experiments. Statistics: Mann-Whitney (d) or paired t-test (b, f), *p≤0.05, **p≤0.01, ***p≤0.001.

Much of our knowledge of the dynamics of leukocyte extravasation comes from the intravital study of neutrophils that are recruited in large numbers at kinetically well-defined times following immune challenge (Nourshargh and Alon, 2014). The relative paucity of CD4^+^ effector T cells entering the inflamed tissue at any given time has hindered the intravital study of their tissue entry. In a mouse model of cerebral malaria, CXCL10 is strongly induced in the brain endothelium upon infection and appears to help promote T cell-endothelial adhesion(Sorensen et al., 2018). In contrast, the blood vessels within the OVA/CFA inflamed dermis did not express detectable levels of BFP-CXCL10 or RFP-CXCL9 by intravital imaging (Fig. 1c, 1h). However, the perivascular localization of BFP-CXCL10^+^ clusters raised the possibility that locally recruited chemokine-producing immune cells may direct targeted T cell extravasation. To test this notion, we i.v. injected fluorescently labeled OVA-specific OT-II TCR transgenic Th1 cells into OVA/CFA-immunized REX3 mice and used IV-MPM to detect the position of incoming Th1 cells relative to the BFP-CXCL10^+^ clusters. As early as 1-2 h post cell transfer, Th1 cells were observed to enter the dermis through vessels adjacent to BFP-CXCL10^+^ clusters (Fig. 3a, b, Video S4, S5). By 18 h post cell transfer, OT-II-Th1 cell accumulation in the dermis was notably clustered at perivascular sites of BFP-CXCL10 expression (Fig. 3c-e; Video S6).

**Figure 3.**
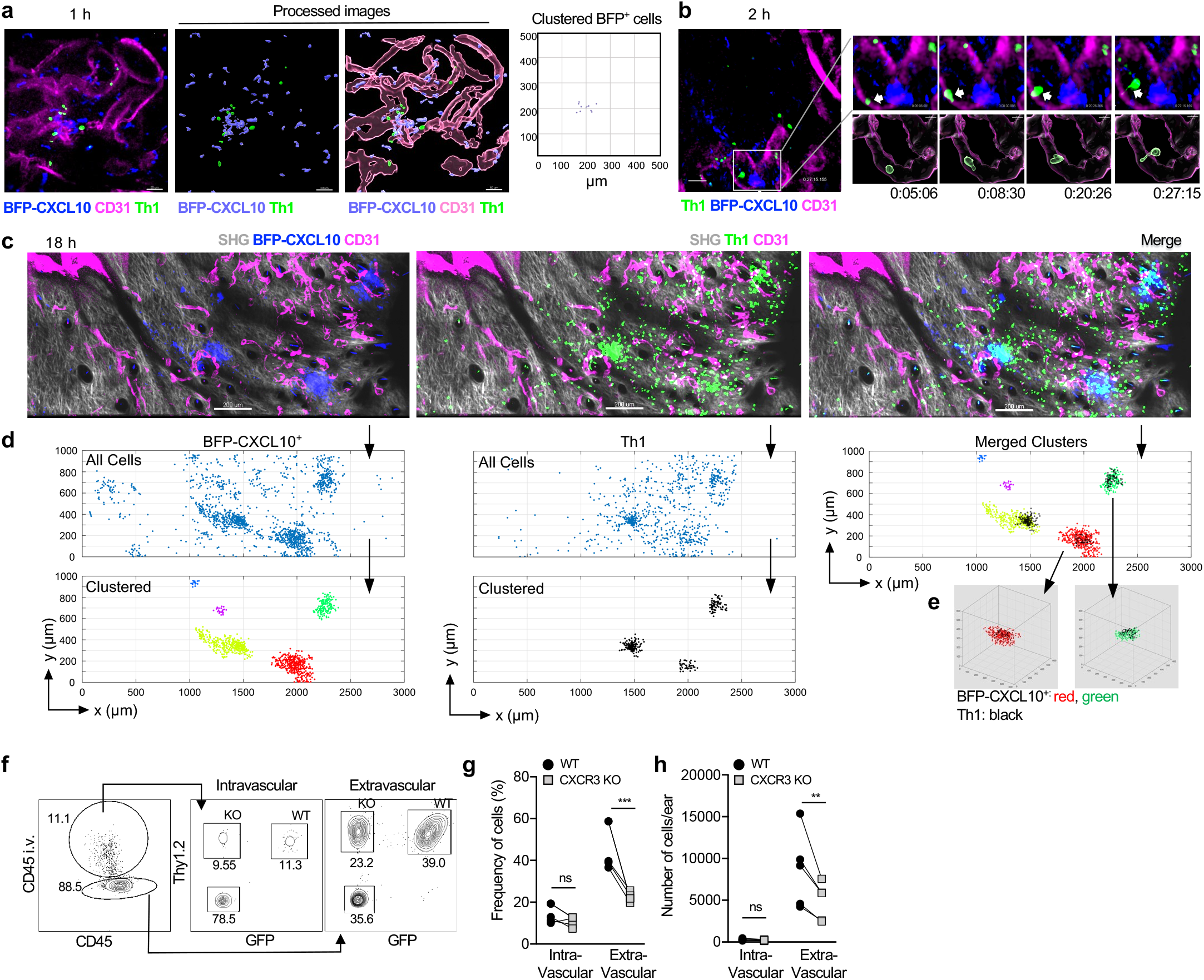
Th1 cells preferentially enter the skin at BFP-CXCL10^+^ clusters in a CXCR3-mediated manner. Fluorescently labeled OT-II Th1 cells were transferred i.v. into d5 OVA/CFA immunized REX3 mice and IV-MPM of the inflamed dermis performed hours after cell transfer. **(a)** 1 h post Th1 cell (green) transfer, CD31^+^ vessel (magenta), BFP-CXCL10^+^ cells (blue). Right panel, BFP-CXCL10^+^ cluster identification using the DBSCAN-based algorithm. **(b)** 2h post Th1 cell (green) transfer, CD31^+^ vessel (magenta), BFP-CXCL10^+^ cells (blue). Insert (upper right): time lapse images of Th1 cell (arrow) transmigrating a CD31^+^ vessel. Processed images (lower right): CD31^+^ vessels reconstructed and volumetrically rendered in 3D using Imaris. **(c)** 18h post Th1 cell (green) Maximal z-projection images: gray, SHG; scale bar 50 μm. Representative of 2 independent experiments, 3 mice, 6 imaging volumes. **(d)** Cluster identification in 3D using DBSCAN-based algorithm of BFP-CXCL10^+^ cells (left panels) and OT-II Th1 cells (middle panels) from (d) (R=50 μm, N=10 neighbors). Top panels, all cells; bottom panels, clustered cells. Merged clusters (top, right panel). **(e)** 3D representation of BFP-CXCL10^+^:T cell co-clusters from (d). **(g-h)** Co-transferred WT and CXCR3 KO OT-II Th1 cells in OVA/CFA immunized Thy1-congenic mice. **(f)** Representative plots, **(g)** frequency and **(h)** number of intravascular (CD45 i.v.) versus extravascular Th1 cells. 3 independent experiments, 3-5 mice per experiment. Stats by paired t-test (g, h), ^ns^ p>0.05, **p≤0.01, ***p≤0.001.

To confirm that the CXCR3-driven Th1 cell accumulation in the perivascular region was indeed a result of local extravasation as opposed to intravascular accumulation, WT and CXCR3 KO OT-II-Th1 cells were co-transferred to OVA/CFA immunized mice and their intravascular versus extravascular location distinguished by acute i.v. injection of anti-CD45 Ab 2 minutes prior to sacrifice followed by flow cytometry(Anderson et al., 2014). The majority of WT Th1 cells were extravascular (not labelled by the i.v. anti-CD45 Ab) (Fig. 3 f-h) consistent with their local exit from the blood into the dermal interstitium at sites of BFP-CXCL10^+^ perivascular clusters. Fewer CXCR3 KO OT-II-Th1 cells were present extravascularly, despite a similar intravascular pool (Fig. 3 f-h), highlighting a role for CXCR3 in tissue entry. Overall, these data suggest that perivascular clusters of CXCL10 production serve as ‘hot spots’ for CXCR3-mediated Th1 tissue entry.

### BFP-CXCL10^+^ clusters enriched for myeloid CD11b^+^CD11c^+^MHC-II^+^ cells

To determine the role of these perivascular clusters in shaping Th1 cell function, we next defined the cellular sources of the chemokines using flow cytometry. Greater than 95% of the CD45^+^ RFP-CXCL9^+^ and BFP-CXCL10^+^ cells were CD11b^+^ myeloid cells and enriched for a subset with MHC-II expression (Fig. 4a, b). Using tSNE analysis of a multiparameter flow cytometry panel aimed at distinguishing major myeloid cell subsets(Guilliams et al., 2016; Nakano et al., 2015) (Fig. S6), the BFP-CXCL10^+^ cells were found to be restricted to a limited number of clusters, one of which was coincident with a subset of the MHC-II^+^ cells (Fig. 4c). Further analysis, defined the predominant BFP-CXCL10^+^ cells to be within monocyte-derived DC (moDC: Ly6G^low^CD64^+^F4/80^+^CD88^+^CD11c^+^MHC-II^+^) and monocyte/macrophage subsets. There was little CXCL9 or CXCL10 expression within the classical DC subsets (cDC) (Fig. 4d-f), predominantly of the cDC2 subset (Fig. S6), or B cells (Fig. 4d-f). Expression levels of BFP extended over a number of logs by flow cytometry, with different innate cell types expressing distinct levels of BFP (Fig. 4f, Fig. S6), but moDCs and MHC-II^+^ mono-macs included sub-populations that had the highest levels of BFP-CXCL10 (Fig. 4f) suggesting that the high chemokine expressing cells are also cells capable of presenting antigen and activating Th1 cells.

**Figure 4.**
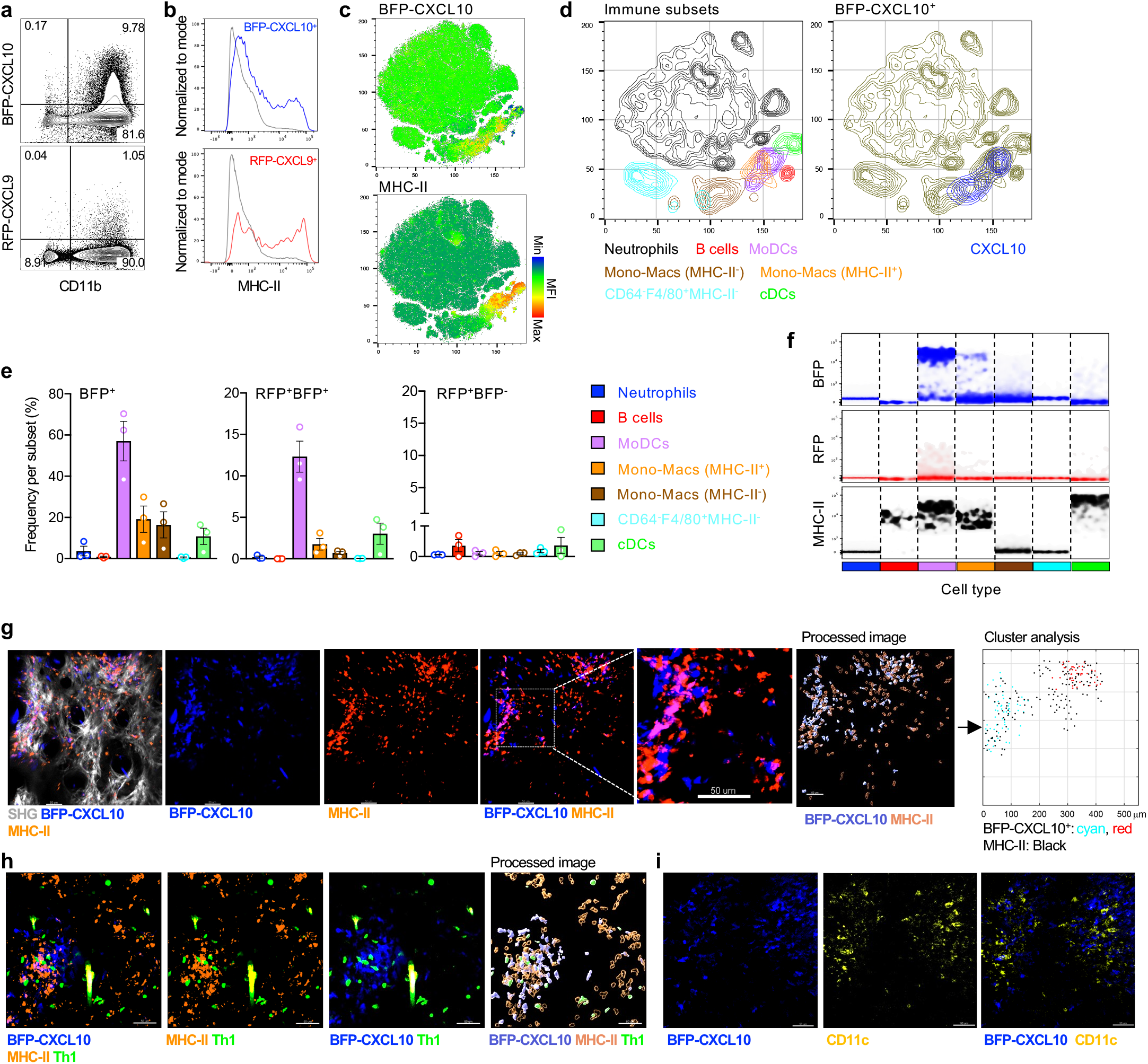
BFP-CXCL10^+^ myeloid cell clusters enriched for CD11c^+^MHC-II^+^ cells. **(a)** Representative dot plots of BFP-CXCL10 and RFP-CXCL9 expression by CD11b^+^ cells within the d5 OVA/CFA immunized REX3 dermis. **(b)** Representative histograms of MHC-II expression by BFP-CXCL10^+^ (blue) and BFP-CXCL9^+^ (red) CD45^+^ cells from (a). Gray: RFP-CXCL9^-^ BFP-CXCL10^-^ CD45^+^ cells. **(c)** tSNE analysis on singlet live CD45^+^ cells using concatenated multi-color flow cytometry data. **(d)** Immune subsets gated by manual flow cytometry analysis displayed on tSNE plots (left panel). Overlay of BFP-CXCL10^+^ cells (right panel). **(e)** Frequency of BFP-CXCL10^+^ cells (left), BFP-CXCL10^+^RFP-CXCL9^+^ cells (middle) and BFP-CXCL10^-^RFP-CXCL9^+^ cells (right) within each of the immune subsets in (d). Bars represent mean ± SEM. 3 independent experiments, 3-4 mice per experiment. **(f)** Density plots for BFP-CXCL10, RFP-CXCL9 and MHC-II expression within each subtype in (e). **(g)** MHC-II^+^ (orange) staining relative to the BFP-CXCL10^+^ cluster and **(h)** colocalization of MHC-II^+^, BFP-CXCL10^+^ and Th1 cells. Right panel, cluster analysis of MHC-II and BFP-CXCL10 cells using the DBSCAN-based algorithm. **(i)** CD11c^+^ (yellow) cells relative to BFP-CXCL10^+^ clusters (blue), d5 OVA/CFA immunized REX3 ears. Representative maximal z-projection images by IV-MPM from 2 independent experiments, 2-3 mice per imaging session, >10 imaging volumes, scale bar 50 µm.

To spatially correlate the cell subsets identified by flow cytometry with the BFP-CXCL10^+^ perivascular clusters in situ in the inflamed dermis, antibodies against MHC-II or CD11c were i.d. injected directly into the immunized REX3 ears 5 days post immunization when the BFP-CXCL10^+^ clusters had formed. Using IV-MPM, we identified dense clusters of MHC-II^+^ cells co-localized with BFP-CXCL10^+^ clusters (Fig. 4g). Moreover, BFP-CXCL10^+^ and MHC-II^+^ co-clusters were preferred sites of Th1 cell accumulation (Fig. 4h). CD11c^+^ cells were also co-localized to BFP-CXCL10^+^ clusters (Fig. 4i). Thus, Th1 cells enter and accumulate within the inflamed tissue in CXCL10^+^ perivascular hubs enriched in both chemokines and APCs.

### Distinct roles for chemokine and antigen in the dynamics of chemokine cluster entry and exit

A pattern of random migration of T cells has been observed within a variety of inflamed tissues including the skin, lung, liver and brain (Egen et al., 2008; Harris et al., 2012; Overstreet et al., 2013). In the absence of evidence for directional migration, the role of chemokines in driving T cell positioning within tissues remains unclear. Data suggest chemokines play roles in micro-anatomical tuning of T cell migration but not in long-range directional chemotaxis. With respect to spatial positioning, interesting studies in zebrafish have raised the possibility that chemokines may also act as stop signals at high concentration in tissues(Sarris et al., 2012). The CXCL9/10 reporter provided a unique opportunity to study the role of CXCR3-chemokines in Th1 tissue positioning in real-time in mice. We used the density-based cluster identification algorithm to define a virtual 3D perimeter of the cluster and then classified T cell tracks according to their interactions with this cluster surface. While the majority of Th1 cells remained inside or outside of the chemokine perivascular cluster over the 60 minute imaging time, co-transferred CXCR3-KO OT-II Th1 cells were much less likely to enter the chemokine-cluster than WT OT-II Th1 cells (Fig. 5a, b). Therefore, in the absence of CXCR3, Th1 cells are not efficiently recruited into the CXCL10-rich cluster. In contrast, there was no difference in the degree of cluster exit between WT and CXCR3-KO OT-II Th1 cells (Fig. 5a, b) suggesting CXCR3-chemokines are not playing a key role in retention of Th1 cells in the CXCL10-rich cluster. Given the antigen-dependency of cluster association (Fig. 2g, Fig. S5), we speculated that cognate antigen would be a major driver in retaining Th1 cells within the CXCL10-rich cluster. As predicted, fewer OT-II Th1 cells were present within the non-cognate antigen KLH/CFA clusters than OVA-containing clusters during the imaging time (Fig. 5c) but OVA-specific Th1 cells were also far more likely to leave the KLH-containing clusters compared to the OVA-containing clusters (Fig. 5c, d). These findings indicate that Th1 cells require CXCR3 chemokine signals to efficiently position within CXCL10-clusters but depend on cognate interactions with antigen-bearing cells to be retained.

**Figure 5.**
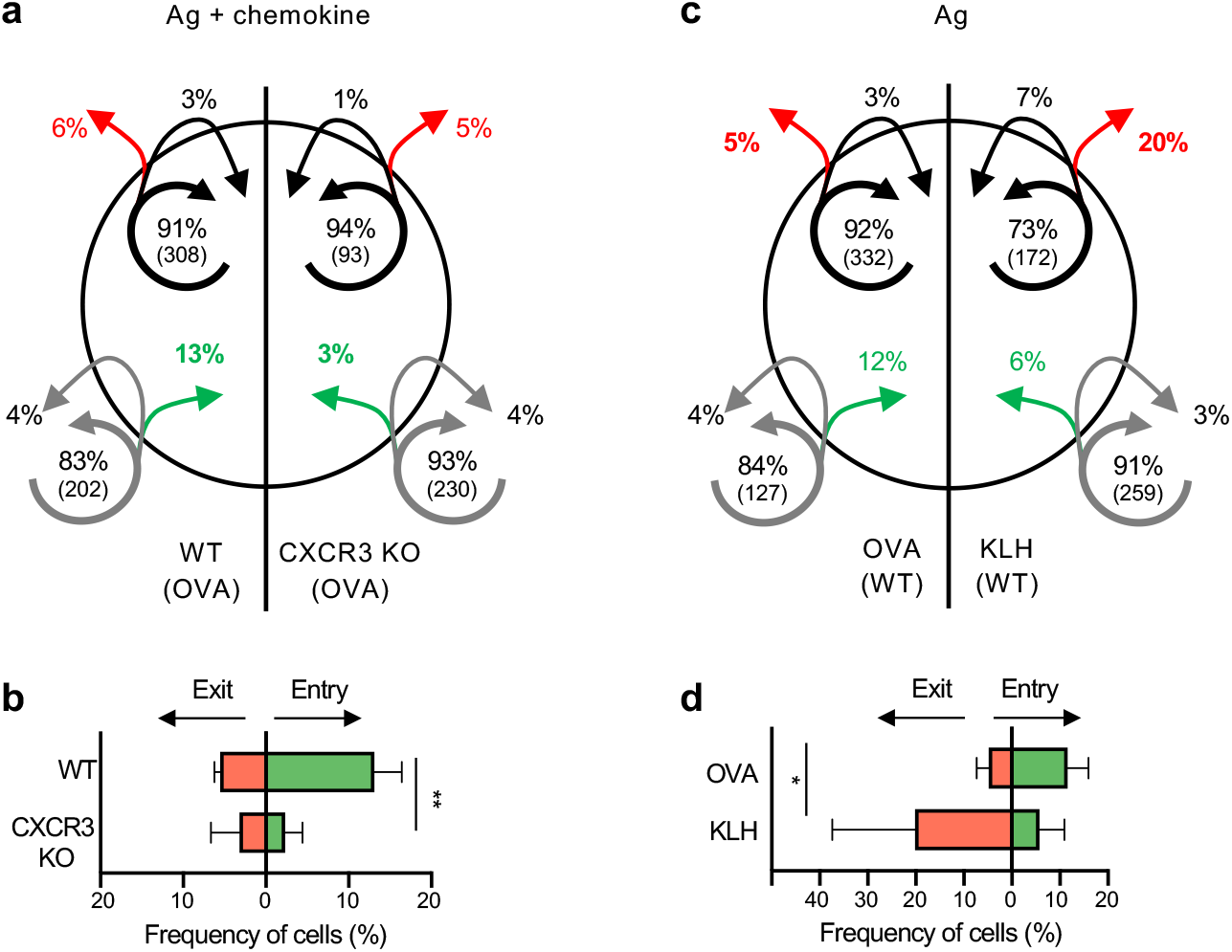
Distinct roles for CXCR3 and cognate antigen in cluster dynamics. IV-MPM analysis of Th1 cell movement in the dermis of REX3 d5 OVA/CFA or KLH/CFA immunized mice. Virtual 3D BFP-CXCL10^+^ clusters in the immunized dermis were defined as in Fig. 1. Transferred OT-II Th1 cell entry and exit from BFP-CXCL10^+^ clusters was tracked over a 60 min imaging period, classified according to their relative location to the cluster perimeter using Python. **(a)** Pattern of movement of co-transferred WT (left) and CXCR3-KO OT-II Th1 (right) cells in the dermis of OVA/CFA immunized mice. The circle represents the BFP-CXCL10^+^ cluster. Arrows indicate the movement of cells relative to the BFP-CXCL10^+^ cluster: green, entered the cluster; red, exited the cluster. Arrow tail position (inside or outside of the BFP-CXCL10^+^ surface) indicates the region in which the track began while arrow head position indicates the region in which the track ended over the 60 min imaging window. Shown are averaged percentages of cells from each genotype that exhibited the same pattern of movement. Numbers in parentheses indicate number of cells in each compartment. **(b)** Frequency of cell entry and exit from the BFP-CXCL10^+^ clusters as in (a). **(c)** Pattern of movement of WT OT-II Th1 cells into and out of BFP-CXCL10^+^ clusters in the dermis of OVA/CFA (left) or KLH/CFA (right), analyzed as in (a). Pooled data from 3 independent experiment, 3-6 mice per group. Bars represent mean + SEM. Stats by two way ANOVA, *p<0.05, ** p< 0.01.

### Perivascular CXCL10^+^ APC clusters optimize antigen-specific T cell activation

To determine the functional impact of CD4^+^ T cell positioning to chemokine-rich regions, we next examined the dynamic behavior of Th1 cells relative to the BFP-CXCL10 clusters. OT-II Th1 cell movement was more confined within the chemokine-rich cluster than outside of the cluster, as measured by the mean squared displacement (Fig. 6a, left panel; Video S7), indicating cues within the cluster that modify Th1 migration. Of note, those CXCR3-deficient Th1 cells that made it into the chemokine-rich cluster were confined in a similar fashion to WT Th1 cells (Fig. 6a, middle panel), thus we find no indication that the CXCR3-chemokines played a role in the location-specific change in Th1 motility. Rather, the presence of cognate antigen dictated the confinement; with no changes in movement of OVA-specific Th1 cells within or outside of KLH-bearing chemokine-rich clusters (Fig. 6a, right panel). Analysis of instantaneous velocity for individual cell tracks relative to the BFP-CXCL10^+^ cluster highlighted remarkable spatiotemporal control of T cell motility (Fig. 6b, c). T cell speed was instantly reduced as the Th1 cell moved from the parenchyma into the CXCL10-rich cluster (Fig. 6b, track 2) and, conversely, Th1 cell speed increased as soon as the cell moved from the cluster out into the interstitium (Fig. 6b, track 3). Further analysis reinforced these observations, Th1 cell motility (average velocity) was reduced within the chemokine-rich cluster compared to movement outside of the chemokine-rich cluster (Fig. 6d, e), independent of CXCR3-chemokines (CXCR3 KO Th1) (Fig. 6e) but dependent on the presence of cognate antigen (OVA versus KLH clusters) (Fig. 6e). Similarly, Th1 cells were more likely to arrest within the chemokine-rich cluster when cognate antigen was present (Fig. 6f) independent of their ability to respond to the CXCR3 chemokines. These data indicate that while CXCR3-chemokines optimize the positioning of the Th1 cells to the chemokine-rich clusters, they are surprisingly not major contributors to Th1 dynamics once within the cluster.

**Figure 6.**
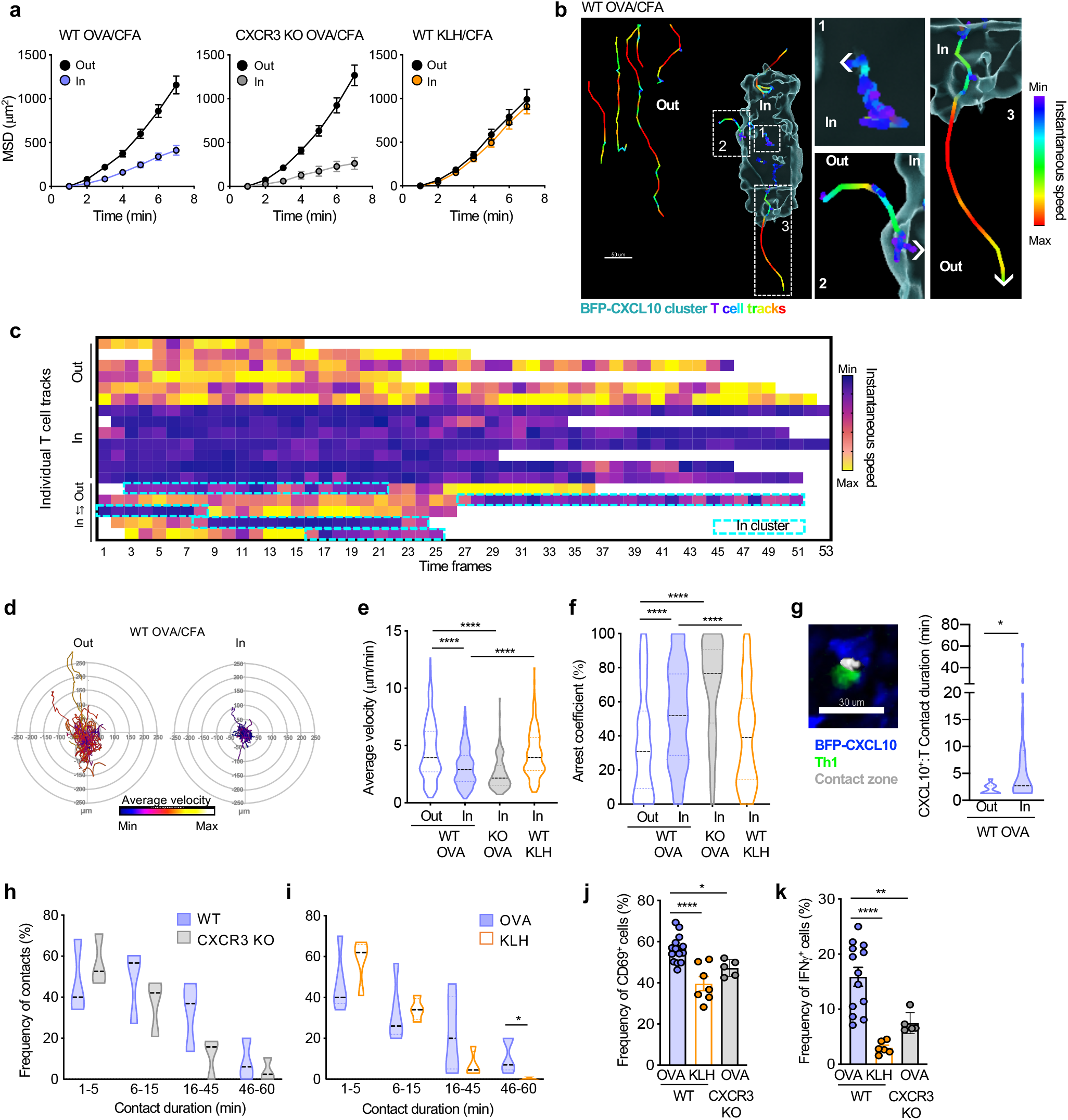
Perivascular BFP-CXCL10^+^ clusters nucleate antigen-specific T:APC interactions to boost function. Motility dynamics of OT-II Th1 cells in the dermis of REX3 d5 OVA/CFA or KLH/CFA immunized mice by IV-MPM. Virtual 3D BFP-CXCL10^+^ clusters were defined by a DBSCAN-based algorithm and Th1 cell motility classified according to their position relative to the cluster perimeter. **(a-d)** T cell motility inside and outside of the BFP-CXCL10^+^ clusters. **(a)** Mean squared displacement (MSD) of WT (left) or CXCR3 KO (middle) OT-II Th1 cells in the OVA/CFA immunized dermis, 30 min IV-MPM. Right panel, WT Th1 cell motility in the OVA/CFA or KLH/CFA immunized dermis. Pooled data from 3 independent experiments, 3-6 mice with 6-12 imaging volumes per group. **(b)** Instantaneous speed of representative tracks of individual Th1 cells in and out of the BFP-CXCL10^+^ cluster, right panels are enlarged regions from the cluster image at left. Cluster surface defined using Imaris surface rendering tool. **(c)** Instantanous speed of individual T cells relative to the BFP-CXCL10^+^ cluster. **(d)** Migratory paths (x-y projections) of OT-II Th1 cells relative to BFP-CXCL10^+^ clusters in the OVA/CFA dermis, 60 min IV-MPM. Representative data from 3 independent experiments, >12 imaging volumes; >30 tracks per plot. **(e)** Average velocity, and **(f)** arrest coefficient, of WT and CXCR3 KO OT-II Th1 cells described in (a). **(g)** Th1:BFP-CXCL10^+^ cell contact time, 60 min IV-MPM. Left, representative IV-MPM image of OT-II Th1 (green):BFP-CXCL10^+^ (blue) cell contacts (white), using the 3D surface overlap rendering tool in Imaris. Right, Th1:BFP-CXCL10^+^ cell contact duration, inside (In) and outside (Out) of BFP-CXCL10^+^ clusters. **(h)** Th1:BFP-CXCL10^+^ cell contact durations for WT and CXCR3 KO Th1 cells, 60 min IV-MPM. **(i)** WT OT-II Th1:BFP-CXCL10^+^ cell contact durations in the OVA/CFA (cognate antigen) and KLH/CFA (non-cognate) immunized dermis. Pooled data from 3 independent experiments, 3-5 mice with 6-10 imaging volumes per group. **(j, k)** Frequency of CD69^+^ **(i)** and IFN-γ^+^ **(j)** WT Th1 cells in the OVA/CFA or KLH/CFA immunized REX3 dermis and CXCR3 KO Th1 cells in the OVA/CFA immunized dermis. Violin plots represent the frequency distribution of the data: black dotted line represents the median and border-colored dotted lines the first and third quartiles. Bars represent mean + SEM. Stats by Mann-Whitney (d-f) or two way ANOVA (g-j), *p<0.05, ** p< 0.01, **** p<0.0001.

We have shown that CXCL10 expression is spatially restricted to perivascular clusters that contain APCs and that Th1 cells have a similar positional bias that is controlled by chemokine-driven localization and antigen-driven retention. Does location to this niche impact Th1 function? To explore the cognate T:APC interactions in these niches, we used an automated 3D surface rendering tool in Imaris (Bitplane) (Fig. 6g)(Gaylo-Moynihan et al., 2019) to calculate the duration of Th1 cell interactions with BFP-CXCL10^+^ APC within and outside of the chemokine-rich clusters. Th1 cells were much more likely to form prolonged contacts with BFP-CXCL10^+^ cells within the perivascular cluster than with BFP-CXCL10^+^ cells dispersed outside of the cluster (Fig. 6g). Consistent with the motility data, it was cognate antigen and not CXCR3-chemokines that impacted stable contacts of more than 45 minutes (Fig. 6h, i). In contrast, functional readouts of the quality of interactions revealed significant contributions from both antigen and CXCR3-chemokines. The upregulation of CD69 as a marker of T cell activation and the frequency of IFNγ producers as a measure of effector function, by ex vivo intracellular cytokine production(McLachlan et al., 2009; Sojka and Fowell, 2011), were predictably compromised in the absence of cognate antigen but were also markedly blunted by the loss of CXCR3 (Fig. 6j, k). Thus, the CXCR3-dependent ability to position to these chemokine-rich perivascular clusters impacts the magnitude of the effector response. We reveal that clustering of CXCL10 chemokine signals and antigen-presenting capacity into specialized niches aids in the targeted recruitment of Th1 cells into tissues and nucleates signals for peripheral T cell activation that prolong T:APC contacts and boost cytokine production. Our data are consistent with the idea that these CXCL10^+^ peripheral activation (PAC-10) niches facilitate initial activation of antigen-specific effector T cells without the need for extensive tissue search.

### An IFNγ-dependent feedback loop enables Th1 cells to orchestrate the composition, number and position of the BFP-CXCL10^+^ clusters

Given the potent action of IFNγ on the upregulation of CXCR3-chemokines, we hypothesized that early recruitment of Th1 cells themselves may establish a positive feedback loop that amplifies the pool of chemokine-producing APCs at the perivascular sites. To test this idea, we enhanced or deleted the pool of CD4^+^ effector cells and determined the effect on CXCL10 expression and cluster formation. Boosting the number of effector T cells by transfer of OTII Th1 cells potently enhanced the number and frequency of CXCL10 single positive (BFP^+^) and CXCL9/10 double positive (BFP^+^RFP^+^) immune cells in the inflamed skin (Fig. 7a), dependent on the presence of cognate antigen (Fig. 7b). Further analysis of the chemokine-expressing immune subsets boosted by Th1 cells, supported a link between the presence of effector T cells and enhanced numbers of MHC-II^+^ CXCL10^+^ moDCs (Fig. 7c) and also revealed enhanced CXCL9 and CXCL10 chemokine expression by both moDCs and cDCs (Fig. 7d). Antibody-mediated blockade of IFNγ (XMG1.2 Ab or control IgG administered 1 day prior to T cell transfer and given daily for 3 days post-T cell transfer) was sufficient to abrogate the Th1 boost in CXCL9 and CXCL10 expression by innate immune subsets (Fig. 7e, f). Similarly, the Th1-mediated induction of MHC-II expression on BFP-CXCL10^+^ cells was lost in the absence of IFNγ (Fig. 7g). These data support a model whereby the interaction of Th1 effectors and APCs in the perivascular clusters results in the cognate upregulation of CXCR3-chemokine expression by APCs (moDCs and cDCs) to amplify the recruitment and activation of incoming Th1 tissue migrants. Consistent with this notion, moDCs and T cells shared a similar kinetic pattern of tissue accumulation, with moDCs numbers increasing coincident with CD4^+^ T cell recruitment between day 3 and day 5 (Fig. 7h, Fig. S7).

**Figure 7.**
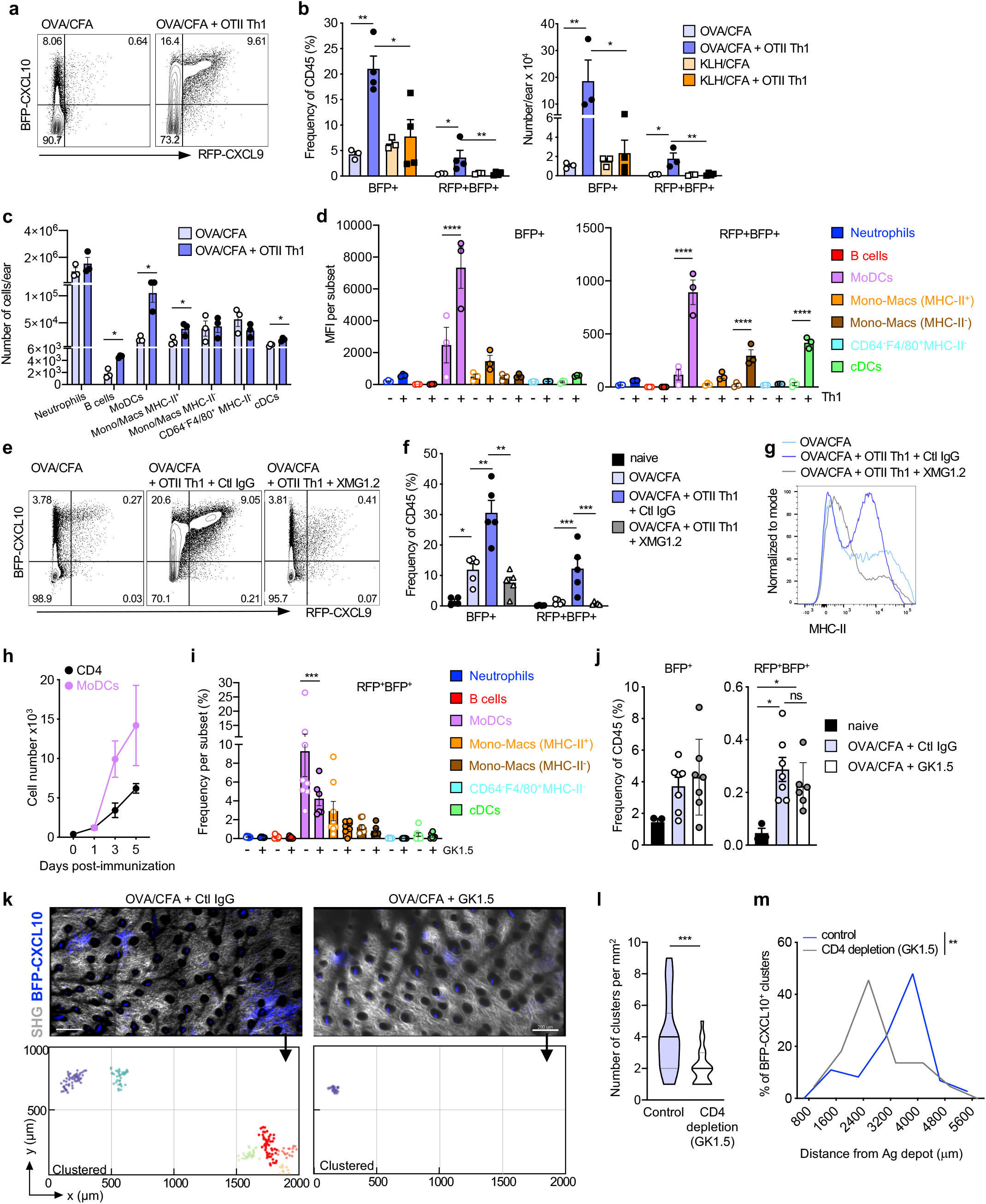
IFNγ-dependent T cell amplification of the chemokine-rich niches. **(a)** Representative plots of BFP-CXCL10 and RFP-CXCL9 expression d5 OVA/CFA immunization ± OT-II Th1 transfer d3 post-immunization. **(b)** Frequency (left) and number (right) of CD45^+^ BFP-CXCL10^+^ and RFP-CXCL9^+^ BFP-CXCL10^+^ cells in mice immunized with OVA/CFA or KLH/CFA ± OT-II Th1 transfer. **(c)** Number and **(d)** BFP-CXCL10 and RFP-CXCL9 MFI within each immune subset. **(a-d)** 3 independent experiments, 3-4 mice per group per experiment. **(e-g)** IFNγ blockade, 0.5 mg XMG1.2 Ab (or control IgG, Ctl IgG) administered i.p. 1 day prior to T cell transfer followed by 3 daily injections of 1 mg. **(e)** Representative plots of BFP-CXCL10 and RFP-CXCL9 expression ± IFNγ-blockade, d5 OVA/CFA. **(f)** Frequency of chemokine expressing cells from (e). **(g)** Representative histograms of MHC-II expression on BFP-CXCL10^+^ cells from (e). **(e-g)** 3 independent experiments, 3-5 mice per group per experiment. **(h)** Number of CD4^+^ T cells and moDCs in the inflamed dermis following OVA/CFA immunization over time. **(i-m)** CD4 depletion, 0.5 mg GK1.5 Ab (or IgG control Ab, Ctl IgG) administered i.p. 1 day prior to OVA/CFA immunization and d1 and d3 post-immunization. **(i)** Frequency of RFP-CXCL9^+^ BFP-CXCL10^+^ within each of the immune subsets ± CD4 depletion. **(j)** Frequency of total BFP-CXCL10^+^ (left) and RFP-CXCL9^+^ BFP-CXCL10^+^ (right) cells, d5 post-immunization. Pooled data from 2 independent experiment, 3-4 mice per experiment. **(k)** Representative dermal tiled images of 8 imaging fields of 512 (x) x 512 (y) x 60 (z) μm from control (left) and CD4 depleted (right) d5 OVA/CFA mice, by IV-MPM, scale bar 200µm. Below, semi-automated 3D cluster reconstruction using DBSCAN-based algorithm. **(l)** Number of identified clusters per mm^2^ of d5 OVA/CFA immunized dermis ± CD4 depletion. **(m)** Frequency distribution of BFP-CXCL10^+^ cell distance from Ag depot ± CD4 depletion. 3 independent experiments, 6 mice, >15 imaging volumes per condition. Bars represent mean + SEM. Statistics by two-way ANOVA with Sidak’s multiple comparisons (b-d, f, I, j) or Mann-Whitney (l, m), *p≤0.05, **p≤0.01, ***p≤0.001, ****p≤0.0001.

To determine the role of CD4^+^ T cells in the establishment and/or maintenance of the PAC-10 niche, we deleted CD4^+^ cells using the anti-CD4 antibody GK1.5 (one day prior to immunization and on days 1 and 3 post-immunization). The loss of CD4^+^ T cells resulted in a marked loss of the MHC-II^+^ CXCL9^+^ CXCL10^+^ moDC population in particular (Fig. 7i), without changes to the overall induction of BFP-CXCL10 and RFP-CXCL9 expression by CD45^+^ cells in the inflamed ear (Fig. 7j). Moreover, the loss of CD4^+^ T cells (and co-dependent moDCs) had a profound impact on the PAC-10 niche (Fig. 7k-m). CD4 depletion reduced the overall number of clusters in the inflamed skin (Fig. 7k, l) and limited their distribution within the tissue (Fig. 7m).

Together, these data reveal the induction of CXCL10^+^ perivascular immune cell clusters that are distributed broadly throughout the inflamed tissue. These PAC-10 niches serve to cluster APC targets at high density in perivascular sites, coupling targeted Th1 recruitment and efficient peripheral activation of Th1 cells. A CD4^+^ T cell positive feedback loop shapes the number and distribution of PAC-10 niches, amplifying the surveillance range of incoming Th1 cells.

## DISCUSSION

Effector CD4^+^ T cells entering inflamed tissues utilize environmental cues to migrate through the tissue interstitium in ‘search’ of cognate ligand. The phenotype, frequency and distribution of a given APC that could support T cell reactivation at inflamed or infected sites is poorly understood. Recent advances in intravital imaging have led to new insight into the molecular machinery utilized by T cells for interstitial migration (Lammermann and Germain, 2014). Yet we remain ill-informed regarding points of T cell tissue entry, the range and modality tissue exploration and the relative positioning of T cell re-activation events (by ligand-bearing APCs) and the infection foci themselves. The findings presented here suggest clustering of chemokine and antigen presentation signals into sub-anatomical structures, close to vessels, optimizes CD4^+^ T cell encounter with APCs in the periphery. This first step in peripheral activation promotes early and selective peripheral activation of antigen-specific effector cells, without the need for extensive tissue search. The use of the CXCL9 and CXCL10 reporter mouse (Groom et al., 2012), enabled the identification of an unrecognized spatial preference for Th1 cell accumulation in the inflamed skin at de-novo organized perivascular sites of CXCL10 expression. Our studies reveal that within the inflamed skin, CXCL9 and CXCL10 expression was enriched in MHC-II expressing cells, predominantly of a moDC phenotype. Mechanistically, colocalization of chemokine-producing APCs and Th1 cells depended on T cell expression of CXCR3 both for site-specific entry into the inflamed tissue and for Th1 positioning to these APC clusters. Constraint of Th1 motility, prolonged APC contacts, and retention within the PAC-10 niches was dependent on recognition of cognate antigen, independent of CXCR3. Nonetheless, chemokine and antigen-derived signals were both critical for the optimization of Th1 effector function within the inflamed skin, chemokines helping to position Th1 cells to these activation niches and antigen helping to keep them there.

Our results provide new insight into the relationship between the point of CD4^+^ T cell tissue entry and the site of initial effector T cell activation in inflamed tissues. Sites of preferred leukocyte entry have been described for neutrophils, where spatially-restricted changes in endothelial expression of adhesins and chemokines, or presence of perivascular macrophages, marks ‘hot-spots’ of neutrophil extravasation(Abtin et al., 2014; Nourshargh and Alon, 2014). Our findings show preferred entry sites for CD4^+^ T cell effectors are shaped by the focal perivascular accumulation of chemokine-producing myeloid cells. We found no evidence for local endothelial expression of CXCL9 and CXCL10 by live imaging, therefore the endothelium likely displays local myeloid-derived chemokines. The ability to now visualize preferred sites of CD4^+^ T cell entry into the inflamed skin will enable important studies to define location-specific changes in the endothelium that support T cell extravasation. Our data are consistent with the idea that the spatial synchronization of vascular exit cues with antigen presentation in the interstitial tissue reduces the scale of T cell search for rare APC targets upon initial tissue entry. This mechanism may also confer an early competitive advantage to those antigen-specific effectors that are recruited along with a host of non-specific effector T cells(Chapman et al., 2005). Therefore, we suggest that PAC-10 niches, at points of tissue entry, provide a rapid screening platform for incoming effector CD4^+^ T cells to promote antigen-driven tissue retention of those effector cells with ‘useful’ specificities. This might be particularly important at early timepoints post-immune challenge when the number of incoming antigen-specific T cells, initial “tissue pioneers”, will be limiting. Moreover, our data support the notion that such effector T cell tissue pioneers activated within the PAC-10 niche, promote the recruitment and/or differentiation of additional chemokine-producing APCs to aid in further Th1 recruitment via an IFNγ-dependent positive feedback loop.

The de novo induction of leukocytic perivascular clusters in the skin has previously been observed at sites of preferred CD8^+^ T cell accumulation in a model of contact hypersensitivity (Natsuaki et al., 2014). This prior study elegantly mapped the sequence of events that established the clusters, with dermal macrophages being critical early initiators of DC and T cell clustering. Here, we reveal a Th1-dependent amplification of the number and tissue-range of these peripheral activation niches. The PAC-10 niches were broadly distributed throughout the inflamed dermis, often 1,000s of microns from the antigen-depot, facilitating a targeted yet tissue-wide response to immune challenge. What determines the location of these perivascular clusters relative to the initial immune challenge remains an interesting unresolved question. Location could be random, driven by the stochastic activation of a critical mass of innate and/or incoming T cells, or directed by the distribution of the infectious agent. Alternatively, these perivascular sites of APC and T cell accumulation may evolve strategically in anatomically-distinct pre-organized niches that could be defined by the steady-state position of resident innate and stromal cells, vasculature and peripheral nerves (Dahlgren and Molofsky, 2019). Emerging data from a variety of tissues has fueled the notion that homeostatic positioning of immune cell types takes advantage of specific micro-anatomically advantageous locations within tissues to optimize the ability to respond to immune challenge. For example, in the LN, naïve CD4^+^ T cells appear to have a peripheral bias in their steady state position at the paracortical region of the LN, perhaps to be better positioned to interact with incoming cDC2 cells (Baptista et al., 2019). Similarly, ILC2s appear to be located within perivascular adventitial cuffs in the lung where colocalized DC and stromal cells (perivascular fibroblasts) are poised to respond to local signals that drain from the tissue parenchyma (Dahlgren et al., 2019) and in niches adjacent to adrenergic neurons in the intestine (Moriyama et al., 2018). Therefore, the ‘scattered’ or broad distribution of PAC-10 niches that we have observed following immunization may reflect similar spatially pre-determined sites poised to respond to immune challenge. In a Th1-dominated challenge these <u>p</u>eripheral <u>ac</u>tivation niches (PAC) are defined by CXCL10, but the signature chemokine may differ depending on the type of immune challenge. Future studies to determine how the position of these early activation niches relate to the subsequent requirement for effector function proximal to the infection foci (Filipe-Santos et al., 2009; Hickman et al., 2015) will be very important. Yet, their broad distribution throughout the inflamed dermis may help to ensure broad initial tissue coverage.

Chemokine receptor deletion approaches clearly highlight a role for chemokines in effector T cell spatial positioning in inflamed tissues; CXCR3-deficient CD8^+^ T cells fail to end up in the right place and hence have reduced ability to clear infections(Hickman et al., 2015; Sung et al., 2012). Mechanistic insight into how the loss of the chemokine sensing results in mis-localization in such *in vivo* models is limited due to the inability to define 3D chemokine gradients in situ. Estimations based on CCL21-producing lymphatic vessels suggest steeply decaying functional gradients from the cellular source, being undetectable for small cells such as lymphocytes within 25 microns of the cellular source(Weber et al., 2013). Thus, the spatially discrete chemokine-rich platforms illuminated by the REX3 chemokine reporter provided a unique opportunity to study chemokine and antigen contributions to CD4^+^ T cell positioning in situ, in the inflamed skin. We show that Th1 effectors utilize CXCR3 to locate to the PAC-10 niche. Functionally, CXCR3-optimized Th1 activation was controlled at two levels: site-specific extravasation at hot-spots of perivascular chemokine production and enhanced interstitial access to these strategically positioned clusters of antigen presentation. In silico modeling of 2D and 3D T cell migration suggests small directional preferences, or “subtle chemotaxis”, enhances the time to target location and could account for the dependency on CXCR3 for cluster entry (Ariotti et al., 2015). Additional behavior previously implicated in chemokine sensing (Sarris and Sixt, 2015), such as a reduction in speed, migratory arrest close to the source and enhanced T:APC interactions, was not observed to be CXCR3-dependent for Th1 cells in the inflamed skin. Instead, cognate antigen drove the confined migratory patterns within the PAC-10 niche, enhanced the duration of T:APC interactions and promoted Th1 retention within the PAC-10 niche. Therefore, we show that CXCR3 chemokines played a very specific role in fine tuning the location of Th1 cells: promoting/guiding access to chemokine-rich APC clusters rather than influencing their dwell time in these regions. Nonetheless, the functional impact of CXCR3-deficiency was profound, having as great an influence on Th1 activation within the tissue as that observed in the absence of cognate antigen. Thus, chemokine-guided positioning is essential for the location of cognate antigen-bearing cells.

The microanatomical activation niche appears to provide additional support for initial peripheral T cell activation. Th1 interactions with CXCL10^+^ APCs within the cluster were prolonged compared to interactions with CXCL10^+^ APCs outside of the cluster. Chemokines have been implicated in stabilizing T:APC interactions (Friedman et al., 2006; Molon et al., 2005), but the enhanced duration of the interactions with CXCL10^+^ APCs in the cluster does not appear to be chemokine driven; as CXCR3 KO Th1 cells were also confined in cognate-antigen clusters and had similar interaction times with CXCL10^+^ cells. Our data does not rule out however that the chemokine/chemokine receptor interactions support a qualitatively different activation signal, independent of T cell dwell time with APC. This may be achieved by locally enriching chemokine or antigen signals, cross talk between neighboring Th1 cells and/or local accumulation or induction of APCs with superior stimulatory potential. The targeted reduction in chemokine-producing moDCs following depletion of CD4^+^ effectors is consistent with the Th1-dependent positive amplification of these clusters.

Many pathogens evade immune cell recognition by modulating the local chemokine milieu, through local inhibition of chemokine production or the deployment of chemokine-decoys(Antonia et al., 2019; Katzman and Fowell, 2008; Mantovani et al., 2006). Therefore, effector function may be enhanced by a sequential two-step process (Ley, 2014) whereby amplification of the frequency or function of incoming antigen-specific T cells (step 1, exemplified here by inducible PAC-10 niches) is kinetically and spatially separated from the pathogen-infection foci (step 2, movement from activation niches to pathogen foci). Indeed, CD8^+^ T cells were shown to require interactions with interstitial DCs after arriving in the influenza-infected lung for effective antiviral immunity at the infected epithelium (McGill et al., 2008). Activation within peripheral activation niches where APC have been shaped by the inflammatory milieu may provide tissue-specific signals that reinforce the Th differentiation program (Katzman and Fowell, 2008; Ley, 2014) or ‘license’ cells for better subsequent search of the tissue to locate the infection foci or enhanced function once at the infection foci (Odoardi et al., 2012). Perivascular activation niches that couple tissue entry with initial tissue activation may quickly expand or arm incoming antigen-specific T cells to better tackle pathogen clearance elsewhere in the target tissue.

Many immune-pathologies are accompanied by perivascular immune infiltrates. Our work suggests that these immune clusters may reflect a conserved mechanism designed to efficiently boost activation of newly tissue-recruited effector T cells. We identify a spatially distinct peripheral activation niche for Th1 cells, PAC-10, that is regulated by the CXCR3-chemokine system. The perivascular positioning of CXCR3 chemokine-producing APCs provides spatial cues that couple Th1 extravasation with Th1 peripheral activation events, effectively reducing the scale of initial T cell search for limited ligand. The identification of CXCL10^+^ peripheral activation sites provides a unique platform to locate and analyze the functional significance of early Th1 re-activation events in inflamed peripheral tissues. These sites may be important therapeutic targets in chronic inflammation, where disruption of the activation niche may mitigate local inflammation.

## ACKNOWLEDGEMENTS

We thank the members of the Fowell lab for helpful discussions on the studies. Jim Miller provided insightful comments on the manuscript. We gratefully acknowledge support from the following agencies: National Institutes of Health Grants P01 AI02851 and R01 AI070826 to DJF; F32 AI138415 to HP; R25 GM064133 to SN; R01 CA204028 to ADL. We acknowledge the support of the Multiphoton and Flow Cytometry core facilities of the University of Rochester Medical Center.

## AUTHOR CONTRIBUTIONS

HP and DJF designed the experiments; HP, SN, AM, AH performed the experiments; HP, NP, AM, SAL, TDM and YRG performed statistical and/or computational analysis; ALM, JRG and ADL provided reagents and advice on experimental design and data interpretation; HP and DJF wrote the manuscript.

## METHODS

### Mice

B6(Cg)-Tyr^c-2J^/J (Albino B6) mice, OVA-specific OT-II TCR transgenic C57BL/6 mice, B6.PL-Thy1^a^/CyJ (B6 Thy1.1) and Cxcr3*^tm1Dgen/J^* (B6 CXCR3 KO) mice were purchased from The Jackson Laboratory. REX3 (Reports the Expression of CXCR3 ligands) B6 mice were provided by A. Luster, Massachusetts General Hospital. WT C57BL/6 mice were purchased from the National Cancer Institute, Bethesda, MD. REX3 mice were crossed to Albino B6 mice. OT-II TCR transgenic mice were crossed to CXCR3 KO mice. Mice were bred and maintained in the pathogen-free animal facility at the University of Rochester. All animal procedures were approved by the Institutional Animal Care and Use Committee of the University of Rochester.

### T cell culture and adoptive transfers

For *in vitro* generation of effector Th1 cells, naïve CD4^+^ T-cells were isolated from lymph nodes and spleens of OT-II TCR transgenic mice using negative selection with a mouse CD4^+^ T Cell Isolation Kit (Miltenyi Biotec). Splenocytes from C57BL/6 mice were T cell depleted by complement lysis of Thy1.2^+^ cells (clone J1J) and irradiated with 2500 rads to use as antigen presenting cells (APCs). Naive CD4^+^ T cells were plated with irradiated APCs, 1µM cognate OVA peptide and with IL-2 (10 U/ml). For skewing to a Th1 phenotype, IL-12 (20 ng/ml; Peprotech), and anti-IL-4 (40 µg/ml; 11B11) were added to the culture. For skewing to a Th2 phenotype, IL-4 (50ng/ml; Peprotech) and anti-IFNγ (50 µg/ml; XMG1.2) were added to the culture. Cells were split in 1:2 ratio three days later and harvested on day five of culture. Cells were labeled with either the fluorescent dye CFSE (carboxyfluorescein succinimidyl ester; Life Technologies) or CellTracker DeepRed dye (Thermo Fisher scientific). Either 7×10^6^ Th1 cells or 1×x10^6^ Th2 cells were adoptively transferred into recipient REX3 or B6 Thy1.1 mice through retro-orbital injection.

### *In-vivo* treatments and reagents

Mice were immunized intradermally (i.d.) in the ear with 10μg OVA protein or KLH protein (Sigma Aldritch) emulsified in complete Freund’s adjuvant (CFA, Sigma Aldritch). Mice were infected i.d. in the ear with 2×10^5^ infectious (PNA-selected) Leishmania major promastigotes (strain WHOM/IR/-/173) in 10ml PBS as previously described (Fowell et al., 1997). For *in-vivo* CD4^+^ T cell depletion, anti-CD4^+^ antibody (0.5 mg per mouse, GK1.5) administered i.p. 1 day prior to immunization and on days 1 and 3 post immunization. For IFNγ blockade, anti-IFNγ antibody (XMG1.2) administered i.p. 1 day prior to Th1 cell adoptive transfer (0.5 mg per mouse) followed by 3 daily injections of 1 mg per mouse. For intravascular immune cell staining, anti-CD45 antibody (3µg per mouse, 30-F11, Biolegend) was injected retro-orbitally 2 min before mice were sacrificed and the ears were removed for cell isolation, *ex-vivo* cytokine staining and flow cytometry analysis as detailed below.

### Flow cytometry

Mice were euthanized, ears were excised, ventral and dorsal sheets separated and each ear placed in 1ml Liberase DL (1 mg/ml in PBS; Sigma Aldrich) with DNase I (50 μg per ear, Sigma Aldrich), minced and incubated with agitation for 30 min at 37 °C. The digested tissues were disrupted by gently pushing the ear through a metal strainer. Insoluble material was removed by filtration and the pellet was washed twice with 2% neonatal calf serum (NCS) in Hank’s balanced-salt solution (HBSS). Single cell suspensions were stained with 1:1000 Ghost dye violet 510 dead cell stain for 405nm excitation (Tonbo biosciences) in PBS for 30 min followed by a wash with 2% NCS HBSS. Next, cells were incubated for 10 min with 1:10 CD16/CD32 Fc block (2.4G2) in FACS buffer (2% NCS/PBS) followed by staining for 30 min at 4°C with fluorochrome-conjugated antibody mix in FACS buffer. Antibodies used include: anti-CD45 BUV395 (1:500, 30-F11, BD Biosciences), anti-CD64 BV605 (1:200, X54-5/7.1, Biolegend), anti-CD11c BV711 (1:450, N418, Biolegend), anti-CD19 BV786 (1:100, 1D3, BD Biosciences), anti-F4/80 PE-Cy5 (1:200, BM8, Invitrogen), anti-CD88 anti-APC (1:800, 20/70, Biolegend), anti-MHC-II APC-ef780 (1:2000, M5/114.15.2, eBioscience), anti-CD26 PE-Cy7 (1:50, H194-112, Biolegend), anti-XCR1 BV650 (1:200, ZET, Biolegend), anti-Ly6G FITC (1:500, 1A8, Biolegend), anti-CD3e Biotin (1:400, 145-2C11, eBioscience) with Streptavidin BV510 (1:1000, Biolegend), anti-CD45 PB (3ug per mouse anti-CD4 BV605 (1:200, RM4-5, Bioleged), anti-CD4 (1:200, RM4-4, Bioleged), anti-IFNγ (1:500, XMG1.2, Invitrogen) anti-Thy-1.2 PE (1:500, 53-2.1, BD biosciences), anti-CD69 PerCP-Cy5.5 (1:500, H1.2F3, BD Biosciences). For ex vivo intracellular cytokine staining, Brefeldin A (1μg/ml) was added to all digestion and wash buffers to block cytokine secretion (Sojka and Fowell, 2011). Cell were washed, filtered and resuspended in FACS buffer with 25 μg DNase I and 20 μl AccuCheck counting beads (Thermo Fisher scientific) per ear. Samples were collected with LSR Fortessa (BD Biosciences) and analyzed with FlowJo Software (Treestar, Ashland, OR). tSNE analysis: tSNE algorithm in FlowJo (learning configuration, Auto, opt-SNE; iterations, 1000; perplexity, 30) was applied on concatenated non-gated or manually gated multi-color flow cytometry data from inflamed REX3 ears.

### Luminex

Ears of mice were excised, digested, and disrupted as above in a total volume of 500µl and kept on ice. Homogenates were centrifuged and the supernatant collected and frozen at −80 C before use. Ear homogenate supernatants were assayed for chemokines and cytokines with the Milliplex MAP Mouse Cytokine/Chemokine Magnetic Bead Panel kit (Millipore). Samples were collected and analyzed using BioPlex 200 reader (BioRad).

### Intravital multiphoton imaging

Mice were anesthetized with isoflurane (induction ∼4%; maintenance ∼1-2%, in air) with an isoflurane vaporizer-ventilation machine (M3000R; Lei Medical). Once mice were anesthetized, the ventral side of the ear pinna was taped to a coverslip. Mice were placed on a custom-made platform for imaging (Gaylo et al., 2016a). Body temperature was maintained with a heated blanket (Kent Scientific) and a heating block (WPI). The microscope objective was heated (Bioptechs) to 36 °C to maintain constant dermal temperature during imaging. Images were acquired with an Olympus FVMPE-RS twin-laser multiphoton system equipped with 4 photomultipliers tubes (PMTs) with blue, green, near-red and far-red filters (Semrock). Fluorescence was collected with an Olympus XLPlanN 25x objective (numerical aperture, 1.05) for deep-tissue multiphoton imaging and was detected with three proprietary photomultipliers. Fluorescence excitation was achieved by a sequential scan of Spectra-Physics DeepSee-MaiTai HP Ti:Sapphire laser tuned to 800 nm and Spectra-Physics InsightX3 laser tuned to 920 nm with DM690-870 and DM690-1050 dichroic mirror. The *z*-stack images (512 × 512 pixels) were 60µm in thickness acquired with a vertical resolution of 2–5 µm and a lateral pixel size of 994 nm. For time-series analyses, three-dimensional stacks were acquired every ∼60s. For acute detection of the MHC-II^+^ and CD11c^+^ cells in the inflamed ear, a 1:1 mixture of anti-CD11c FITC (N418, eBioscience) or anti-MHC-II AF647 antibody (M5/114.15.2, eBioscience) with anti-CD16/32 antibody (Fc Block, 2.4G2) were dialyzed against PBS to remove sodium azide. Immunized mice were injected i.d. in the ear with the antibody mix 2 hours prior to IV-MPM imaging. Blood and lymphatic vessels were stained with anti-CD31 AF647 (10μg per mouse, retro-orbital 15 min prior to imaging, clone 390, Biolegend) and anti-LYVE-1 AF647 (10μg per mouse, i.d. 2 hours prior to imaging) respectively.

### Analysis of multiphoton data using Imaris and Python

Raw imaging data were processed with Imaris software (Bitplane). A 3×3 median filter was used to diminish noise and T cells were tracked with automated algorithms with manual correction. Cell tracks lasting for less than 8 min were excluded from analyses. No minimum cell-displacement criteria were imposed that would have excluded non-motile cells. Mean squared displacement (MSD) was calculated from raw imaging data using a custom Python script. The MSD was calculated from trajectory data using the following equation: 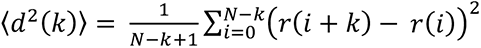 where *N* is the total length of the data, *r* is the trajectory vector, and *k* is the normalized time delay, which is the elapsed time divided by the time per step. Average velocity and instantaneous speed were calculated in Imaris using track speed mean and speed formulas, respectively. Arrest coefficients were calculated as the ratio between the time that a cell was not moving (instantaneous speed <2 μm/min) and the total time the cell was observed. Cell tracks were linearly interpolated in order to maintain consistent time windows for analysis. All analysis was performed through custom-written Python code available upon request. Movies were created in Imaris, with animation and titles added with Adobe Primer Pro.

#### T:CXCL10^+^ cell contacts

cell:cell interactions were detected within the 3D volume reconstituted from all z stacks (over time). T cells and BFP-CXCL10^+^ cells were reconstructed and volumetrically rendered in 3D using Imaris. We first binary masked the signal from each channel (threshold for masking is based on object size and fluorescence intensity). A separate channel was created based on overlapped pixels (threshold is absolute intensity). These overlapped pixels subjected to surface rendering in 3D to represent and visualize T cell-CXCL10^+^ cell contacts. To measure duration of the contacts, 3D surfaces were tracked over time.

#### Relative cell distance

CXCL10^+^ cell distance from blood vessels or T cell distance from CXCL10^+^ cells: vessels and cells were volumetrically rendered in 3D using Imaris. Distance transformation Xtension was applied to the CD31^+^ or CXCL10^+^ surface, resulted in a new channel with the intensities equal to the distance from the surface object. Minimal distance of each CXCL10^+^ cell from the closest CD31^+^ vessel, or T cell from the closest CXCL10^+^ cell, was calculated using Imaris.

#### 3D CXCL10^+^ cluster reconstruction and T cell dynamics

Clusters of CXCL10^+^ cells were defined and reconstituted in 3D using a Python code that utilizes the unsupervised machine learning algorithm DBSCAN (Density-based spatial clustering of applications with noise)(Ester et al., 1996). There are two parameters to the algorithm; maximum distance between points, radius (R) and minimum number of neighbors (N) within R to define as part of the cluster. Density cutoffs of R=50 µm and N=10 neighbors were determined by grid-search where combinations of different R and N were plotted to find inflection points after which return on metric of choice is incremental. Cells outside the identified clusters were treated as noise and therefore ignored in cluster analysis. To determine T cell dynamics relative to the identified clusters (entry, exit. re-entry etc.), T cells were registered at the start of the imaging session and thereafter per time point, as positioned inside or outside of the identified clusters. For analysis of T cell motility inside versus outside of clusters, only cells that positioned the entire imaging session inside clusters compared with cells that spent the entire imaging session outside of the clusters, were included. All analysis was performed through custom-written Python code available upon request.

## QUATIFICATION AND SATISTICAL ANALYSIS

GraphPad Prism was used for all statistical tests. Most analyses used the nonparametric Mann-Whitney or Kolmogorov-Smirnov tests to compare 2 treatment groups. ANOVA tests were used across multiple groups, with multiple comparisons. * = p ≤0.05, ** = p ≤0.01, *** = p ≤0.001, **** = p <0.0001, ns = p > 0.05.

## SUPPLEMENTAL VIDEO DESCRIPTIONS

### Video S1. Perivascular BFP-CXCL10^+^ clusters

REX3 mice were immunized with OVA protein emulsified in CFA in the ear pinna and the inflamed dermis imaged by IV-MPM five days later. Distribution of BFP-CXCL10^+^ cells (blue) and CD31^+^ vessels (red) in the dermis. The video represents a 2D maximal z-projection time series of a 60μm-thick imaging volume; scale bar 50µm.

### Video S2. CXCR3-dependent Th1 cell accumulation in BFP-CXCL10^+^ clusters

CFSE-fluorescently labeled WT and DeepRed-fluorescently labeled CXCR3 KO OT-II Th1 cells were transferred i.v. into d2 REX3 mice immunized with OVA/CFA and IV-MPM of the inflamed dermis performed three days later. Distribution of WT (green) and CXCR3 KO (magenta) OT-II Th1 cells relative to BFP-CXCL10^+^ clusters (blue). The video represents a 2D maximal z-projection time series of a 60μm-thick imaging volume; scale bar 50µm.

### Video S3. Antigen-dependent Th1 cell accumulation in BFP-CXCL10^+^ clusters

CFSE-fluorescently labeled OT-II Th1 cells were transferred i.v. into d2 REX3 mice immunized with OVA/CFA (cognate antigen) or KLH/CFA (non-cognate antigen) and IV-MPM of the inflamed dermis performed three days later. Side by side distribution of OT-II Th1 cells (green) and BFP-CXCL10^+^ cells (blue) in the presence of non-cognate antigen (left) and cognate antigen (right). The video represents a 2D maximal z-projection time series of a 60μm-thick imaging volume; scale bar 50µm.

### Video S4. Th1 cell skin entry adjacent to BFP-CXCL10^+^ cluster

CFSE-fluorescently labeled OT-II Th1 cells were transferred i.v. into d5 OVA/CFA immunized REX3 mice and IV-MPM of the inflamed dermis performed 2h post Th1 cell (green) transfer. Th1 cell (arrow) transmigrating a CD31^+^ vessel (magenta) next to BFP-CXCL10^+^ cluster (blue). Processed images: CD31^+^ vessels (pink) and OT-II Th1 cell (green) reconstructed and volumetrically rendered in 3D using Imaris. The video represents a 2D maximal z-projection time series of a 60μm-thick imaging volume; Scale bar 50µm.

### Video S5. Th1 cells crawling in vessels adjacent to BFP-CXCL10^+^ cluster

CFSE-fluorescently labeled OT-II Th1 cells were transferred i.v. into d5 OVA/CFA immunized REX3 mice and IV-MPM of the inflamed dermis performed 3h post Th1 cell (green) transfer. Th1 cells crawling in CD31^+^ vessels (red) adjacent to BFP-CXCL10^+^ cluster (blue); tissue collagen matrix visualized by SHG (gray). Processed images: CD31^+^ vessels (pink) and OT-II Th1 cell (green) reconstructed and volumetrically rendered in 3D using Imaris. The video represents a 2D maximal z-projection or rotated 3D volume time series of a 60μm-thick imaging volume; Scale bar 50µm.

### Video S6. Th1 cell accumulation in BFP-CXCL10^+^ clusters

CFSE-fluorescently labeled OT-II Th1 cells were transferred i.v. into d5 OVA/CFA immunized REX3 mice and IV-MPM of the inflamed dermis performed 18h post Th1 cell transfer. Cluster identification in 3D using DBSCAN-based algorithm in Python (R=50 μm, N=10 neighbors) of BFP-CXCL10^+^ cells (multi-color) and OT-II Th1 cells (black). Still plot represent a 2D maximal z-projection of a 60μm-thick tiled imaging volume; Rotating plots represent each a single BFP-CXCL10^+^:T cell co-cluster in 3D; Axes represent distance in µm.

### Video S7. Th1 cell confinement within BFP-CXCL10^+^ cluster

CFSE-fluorescently labeled WT OT-II Th1 cells were transferred i.v. into d2 REX3 mice immunized with OVA/CFA and IV-MPM of the inflamed dermis performed three days later. Distribution of WT OT-II Th1 cells (green) relative to BFP-CXCL10^+^ cluster (blue); tissue collagen matrix visualized by SHG (gray). Processed video: OT-II Th1 cells (green) reconstructed and volumetrically rendered in 3D using Imaris. T cell trajectories via 3D surface creation in Imaris displayed in dragon-tail mode; tracks color coded by average velocity (track speed mean) in Imaris. The video represents a 2D maximal z-projection time series of a 60μm-thick imaging volume; scale bar 50µm.

## REFERENCES

Abtin, A., Jain, R., Mitchell, A.J., Roediger, B., Brzoska, A.J., Tikoo, S., Cheng, Q., Ng, L.G., Cavanagh, L.L., von Andrian, U.H., et al. (2014). Perivascular macrophages mediate neutrophil recruitment during bacterial skin infection. Nat Immunol 15, 45–53.

Anderson, K.G., Mayer-Barber, K., Sung, H., Beura, L., James, B.R., Taylor, J.J., Qunaj, L., Griffith, T.S., Vezys, V., Barber, D.L., and Masopust, D. (2014). Intravascular staining for discrimination of vascular and tissue leukocytes. Nat Protoc 9, 209–222.

Antonia, A.L., Gibbs, K.D., Trahair, E.D., Pittman, K.J., Martin, A.T., Schott, B.H., Smith, J.S., Rajagopal, S., Thompson, J.W., Reinhardt, R.L., and Ko, D.C. (2019). Pathogen Evasion of Chemokine Response Through Suppression of CXCL10. Front Cell Infect Microbiol 9, 280.

Ariotti, S., Beltman, J.B., Borsje, R., Hoekstra, M.E., Halford, W.P., Haanen, J.B., de Boer, R.J., and Schumacher, T.N. (2015). Subtle CXCR3-Dependent Chemotaxis of CTLs within Infected Tissue Allows Efficient Target Localization. J Immunol 195, 5285–5295.

Baptista, A.P., Gola, A., Huang, Y., Milanez-Almeida, P., Torabi-Parizi, P., Urban, J.F., Jr., Shapiro, V.S., Gerner, M.Y., and Germain, R.N. (2019). The Chemoattractant Receptor Ebi2 Drives Intranodal Naive CD4(+) T Cell Peripheralization to Promote Effective Adaptive Immunity. Immunity 50, 1188–1201 e1186.

Chapman, T.J., Castrucci, M.R., Padrick, R.C., Bradley, L.M., and Topham, D.J. (2005). Antigen-specific and non-specific CD4+ T cell recruitment and proliferation during influenza infection. Virology 340, 296–306.

Dahlgren, M.W., Jones, S.W., Cautivo, K.M., Dubinin, A., Ortiz-Carpena, J.F., Farhat, S., Yu, K.S., Lee, K., Wang, C., Molofsky, A.V., et al. (2019). Adventitial Stromal Cells Define Group 2 Innate Lymphoid Cell Tissue Niches. Immunity 50, 707–722 e706.

Dahlgren, M.W., and Molofsky, A.B. (2019). Adventitial Cuffs: Regional Hubs for Tissue Immunity. Trends Immunol 40, 877–887.

Egen, J.G., Rothfuchs, A.G., Feng, C.G., Horwitz, M.A., Sher, A., and Germain, R.N. (2011). Intravital imaging reveals limited antigen presentation and T cell effector function in mycobacterial granulomas. Immunity 34, 807–819.

Egen, J.G., Rothfuchs, A.G., Feng, C.G., Winter, N., Sher, A., and Germain, R.N. (2008). Macrophage and T cell dynamics during the development and disintegration of mycobacterial granulomas. Immunity 28, 271–284.

Ester, M., Kriegel, H.-P., Sander, J., and Xu, X. (1996). A Density-Based Algorithm for Discovering Clusters in Large Spatial Databases with Noise. KDD-96 Proceedings, 226–231.

Filipe-Santos, O., Pescher, P., Breart, B., Lippuner, C., Aebischer, T., Glaichenhaus, N., Spath, G.F., and Bousso, P. (2009). A dynamic map of antigen recognition by CD4 T cells at the site of Leishmania major infection. Cell Host Microbe 6, 23–33.

Fowell, D.J., Magram, J., Turck, C.W., Killeen, N., and Locksley, R.M. (1997). Impaired Th2 subset development in the absence of CD4. Immunity 6, 559–569.

Friedman, R.S., Jacobelli, J., and Krummel, M.F. (2006). Surface-bound chemokines capture and prime T cells for synapse formation. Nat Immunol 7, 1101–1108.

Gaylo, A., Overstreet, M.G., and Fowell, D.J. (2016a). Imaging CD4 T Cell Interstitial Migration in the Inflamed Dermis. J Vis Exp, e53585.

Gaylo, A., Schrock, D.C., Fernandes, N.R., and Fowell, D.J. (2016b). T Cell Interstitial Migration: Motility Cues from the Inflamed Tissue for Micro- and Macro-Positioning. Front Immunol 7, 428.

Gaylo-Moynihan, A., Prizant, H., Popovic, M., Fernandes, N.R.J., Anderson, C.S., Chiou, K.K., Bell, H., Schrock, D.C., Schumacher, J., Capece, T., et al. (2019). Programming of Distinct Chemokine-Dependent and -Independent Search Strategies for Th1 and Th2 Cells Optimizes Function at Inflamed Sites. Immunity 51, 298–309 e296.

Griffith, J.W., Sokol, C.L., and Luster, A.D. (2014). Chemokines and chemokine receptors: positioning cells for host defense and immunity. Annu Rev Immunol 32, 659–702.

Groom, J.R., Richmond, J., Murooka, T.T., Sorensen, E.W., Sung, J.H., Bankert, K., von Andrian, U.H., Moon, J.J., Mempel, T.R., and Luster, A.D. (2012). CXCR3 chemokine receptor-ligand interactions in the lymph node optimize CD4+ T helper 1 cell differentiation. Immunity 37, 1091–1103.

Guilliams, M., Dutertre, C.A., Scott, C.L., McGovern, N., Sichien, D., Chakarov, S., Van Gassen, S., Chen, J., Poidinger, M., De Prijck, S., et al. (2016). Unsupervised High-Dimensional Analysis Aligns Dendritic Cells across Tissues and Species. Immunity 45, 669–684.

Harris, T.H., Banigan, E.J., Christian, D.A., Konradt, C., Tait Wojno, E.D., Norose, K., Wilson, E.H., John, B., Weninger, W., Luster, A.D., et al. (2012). Generalized Levy walks and the role of chemokines in migration of effector CD8+ T cells. Nature 486, 545–548.

Hickman, H.D., Reynoso, G.V., Ngudiankama, B.F., Cush, S.S., Gibbs, J., Bennink, J.R., and Yewdell, J.W. (2015). CXCR3 chemokine receptor enables local CD8(+) T cell migration for the destruction of virus-infected cells. Immunity 42, 524–537.

Iijima, N., and Iwasaki, A. (2014). T cell memory. A local macrophage chemokine network sustains protective tissue-resident memory CD4 T cells. Science 346, 93–98.

Katzman, S.D., and Fowell, D.J. (2008). Pathogen-imposed skewing of mouse chemokine and cytokine expression at the infected tissue site. J Clin Invest 118, 801–811.

Krummel, M.F., Bartumeus, F., and Gerard, A. (2016). T cell migration, search strategies and mechanisms. Nat Rev Immunol 16, 193–201.

Lammermann, T., and Germain, R.N. (2014). The multiple faces of leukocyte interstitial migration. Semin Immunopathol 36, 227–251.

Lammermann, T., and Kastenmuller, W. (2019). Concepts of GPCR-controlled navigation in the immune system. Immunol Rev 289, 205–231.

Ley, K. (2014). The second touch hypothesis: T cell activation, homing and polarization. F1000Res 3, 37.

Mandl, J.N., Torabi-Parizi, P., and Germain, R.N. (2014). Visualization and dynamic analysis of host-pathogen interactions. Curr Opin Immunol 29C, 8–15.

Mantovani, A., Bonecchi, R., and Locati, M. (2006). Tuning inflammation and immunity by chemokine sequestration: decoys and more. Nat Rev Immunol 6, 907–918.

Masopust, D., and Schenkel, J.M. (2013). The integration of T cell migration, differentiation and function. Nat Rev Immunol 13, 309–320.

McGill, J., Van Rooijen, N., and Legge, K.L. (2008). Protective influenza-specific CD8 T cell responses require interactions with dendritic cells in the lungs. J Exp Med 205, 1635–1646.

McLachlan, J.B., Catron, D.M., Moon, J.J., and Jenkins, M.K. (2009). Dendritic cell antigen presentation drives simultaneous cytokine production by effector and regulatory T cells in inflamed skin. Immunity 30, 277–288.

Molon, B., Gri, G., Bettella, M., Gomez-Mouton, C., Lanzavecchia, A., Martinez, A.C., Manes, S., and Viola, A. (2005). T cell costimulation by chemokine receptors. Nat Immunol 6, 465–471.

Moriyama, S., Brestoff, J.R., Flamar, A.L., Moeller, J.B., Klose, C.S.N., Rankin, L.C., Yudanin, N.A., Monticelli, L.A., Putzel, G.G., Rodewald, H.R., and Artis, D. (2018). beta2-adrenergic receptor-mediated negative regulation of group 2 innate lymphoid cell responses. Science 359, 1056–1061.

Mrass, P., Takano, H., Ng, L.G., Daxini, S., Lasaro, M.O., Iparraguirre, A., Cavanagh, L.L., von Andrian, U.H., Ertl, H.C., Haydon, P.G., and Weninger, W. (2006). Random migration precedes stable target cell interactions of tumor-infiltrating T cells. J Exp Med 203, 2749–2761.

Nakano, H., Moran, T.P., Nakano, K., Gerrish, K.E., Bortner, C.D., and Cook, D.N. (2015). Complement receptor C5aR1/CD88 and dipeptidyl peptidase-4/CD26 define distinct hematopoietic lineages of dendritic cells. J Immunol 194, 3808–3819.

Natsuaki, Y., Egawa, G., Nakamizo, S., Ono, S., Hanakawa, S., Okada, T., Kusuba, N., Otsuka, A., Kitoh, A., Honda, T., et al. (2014). Perivascular leukocyte clusters are essential for efficient activation of effector T cells in the skin. Nat Immunol 15, 1064–1069.

Nourshargh, S., and Alon, R. (2014). Leukocyte migration into inflamed tissues. Immunity 41, 694–707.

Odoardi, F., Sie, C., Streyl, K., Ulaganathan, V.K., Schlager, C., Lodygin, D., Heckelsmiller, K., Nietfeld, W., Ellwart, J., Klinkert, W.E., et al. (2012). T cells become licensed in the lung to enter the central nervous system. Nature 488, 675–679.

Overstreet, M.G., Gaylo, A., Angermann, B.R., Hughson, A., Hyun, Y.M., Lambert, K., Acharya, M., Billroth-Maclurg, A.C., Rosenberg, A.F., Topham, D.J., et al. (2013). Inflammation-induced interstitial migration of effector CD4(+) T cells is dependent on integrin alphaV. Nat Immunol 14, 949–958.

Sallusto, F., and Lanzavecchia, A. (2000). Understanding dendritic cell and T-lymphocyte traffic through the analysis of chemokine receptor expression. Immunol Rev 177, 134–140.

Sarris, M., Masson, J.B., Maurin, D., Van der Aa, L.M., Boudinot, P., Lortat-Jacob, H., and Herbomel, P. (2012). Inflammatory chemokines direct and restrict leukocyte migration within live tissues as glycan-bound gradients. Curr Biol 22, 2375–2382.

Sarris, M., and Sixt, M. (2015). Navigating in tissue mazes: chemoattractant interpretation in complex environments. Curr Opin Cell Biol 36, 93–102.

Schumann, K., Lammermann, T., Bruckner, M., Legler, D.F., Polleux, J., Spatz, J.P., Schuler, G., Forster, R., Lutz, M.B., Sorokin, L., and Sixt, M. (2010). Immobilized chemokine fields and soluble chemokine gradients cooperatively shape migration patterns of dendritic cells. Immunity 32, 703–713.

Sojka, D.K., and Fowell, D.J. (2011). Regulatory T cells inhibit acute IFN-gamma synthesis without blocking T-helper cell type 1 (Th1) differentiation via a compartmentalized requirement for IL-10. Proc Natl Acad Sci U S A 108, 18336–18341.

Sorensen, E.W., Lian, J., Ozga, A.J., Miyabe, Y., Ji, S.W., Bromley, S.K., Mempel, T.R., and Luster, A.D. (2018). CXCL10 stabilizes T cell-brain endothelial cell adhesion leading to the induction of cerebral malaria. JCI Insight 3.

Sorokin, L. (2010). The impact of the extracellular matrix on inflammation. Nat Rev Immunol 10, 712–723.

Sung, J.H., Zhang, H., Moseman, E.A., Alvarez, D., Iannacone, M., Henrickson, S.E., de la Torre, J.C., Groom, J.R., Luster, A.D., and von Andrian, U.H. (2012). Chemokine guidance of central memory T cells is critical for antiviral recall responses in lymph nodes. Cell 150, 1249–1263.

Veres, T.Z., Kopcsanyi, T., van Panhuys, N., Gerner, M.Y., Liu, Z., Rantakari, P., Dunkel, J., Miyasaka, M., Salmi, M., Jalkanen, S., and Germain, R.N. (2017). Allergen-Induced CD4+ T Cell Cytokine Production within Airway Mucosal Dendritic Cell-T Cell Clusters Drives the Local Recruitment of Myeloid Effector Cells. J Immunol 198, 895–907.

Weber, M., Hauschild, R., Schwarz, J., Moussion, C., de Vries, I., Legler, D.F., Luther, S.A., Bollenbach, T., and Sixt, M. (2013). Interstitial dendritic cell guidance by haptotactic chemokine gradients. Science 339, 328–332.

