## Supplemental Figures for "CXCL10^+^ peripheral activation niches couple preferred sites of Th1 entry with optimal APC encounter"

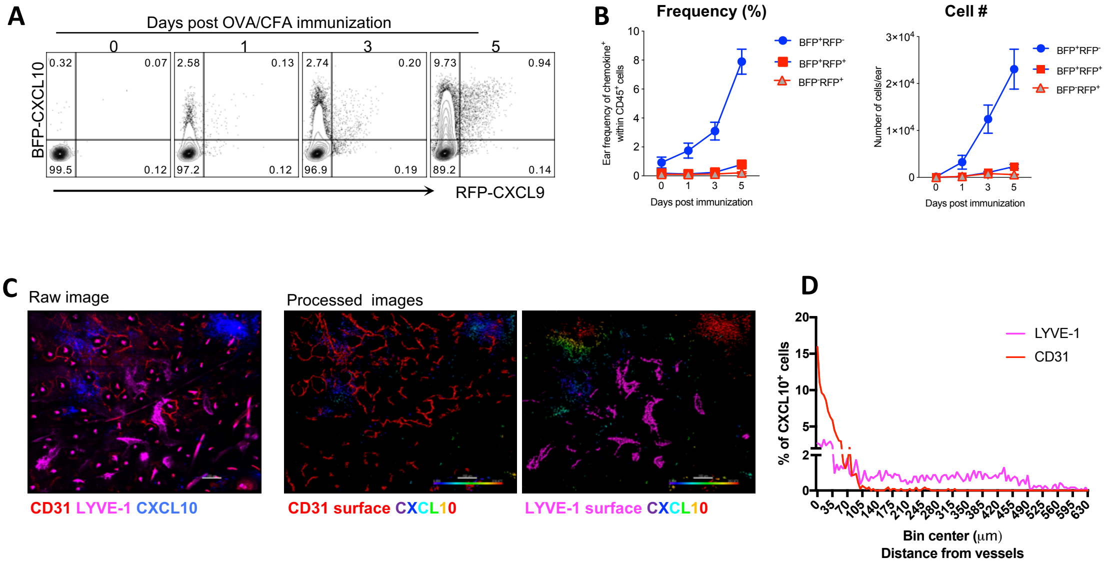


**Figure S1. CXCR3 chemokine expression and distribution within the inflamed skin**

REX3 mice immunized with OVA/CFA in the ear dermis. **(A)** Representative plots and quantification **(B)** of frequency (left) and number (right) of BFP-CXCL10 and RFP-CXCL9 expressing cells within live singlets CD45$^{+}$ cells, by flow cytometry. **(C)** Representative IV-MPM tiled image and **(D)** quantification of BFP-CXCL10$^{+}$ cells distance from CD31$^{+}$ and LYVE-1$^{+}$ vessels within the same representative imaging volume. Scale bar, 100 µm. Anti-CD31 and anti-LYVE-1 antibodies co-injected i.d. to the ear 2 hours prior to imaging.

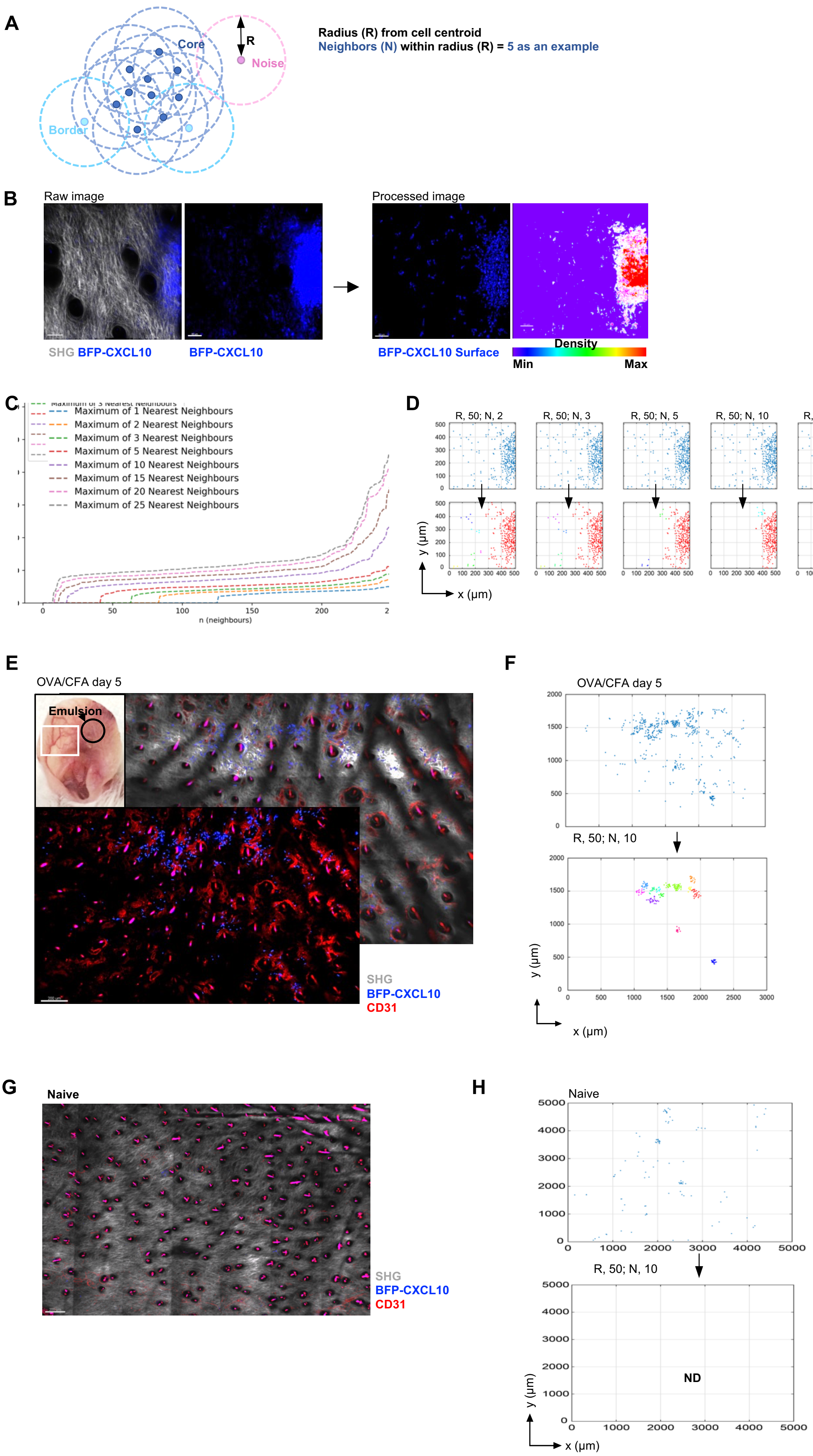

A
Core
Noise
Border
R
Radius (R) from cell centroid
Neighbors (N) within radius (R) = 5 as an example
B
Raw image
Processed image
SHG BFP-CXCL10
BFP-CXCL10
BFP-CXCL10 Surface
Density
Min
Max
C
Maximum of 1 Nearest Neighbours
Maximum of 2 Nearest Neighbours
Maximum of 3 Nearest Neighbours
Maximum of 5 Nearest Neighbours
Maximum of 10 Nearest Neighbours
Maximum of 15 Nearest Neighbours
Maximum of 20 Nearest Neighbours
Maximum of 25 Nearest Neighbours
n (neighbours)
D
R, 50; N, 2
R, 50; N, 3
R, 50; N, 5
R, 50; N, 10
R, 50; N, 15
R, 50; N, 20
All cells
Clusters
y (µm)
x (µm)
E
OVA/CFA day 5
Emulsion
SHG
BFP-CXCL10
CD31
F
OVA/CFA day 5
R, 50; N, 10
y (µm)
x (µm)
G
Naive
SHG
BFP-CXCL10
CD31
H
Naive
R, 50; N, 10
ND
y (µm)
x (µm)

**Figure S2. Selecting criteria for BFP-CXCL10$^+$ cell clustering**

DBSCAN-based machine learning algorithm was developed to unbiasedly and semi-automatically define cell clusters in 3D across different inflamed ears and conditions, using Python. **(A)** Clusters were defined as regions of high density. Each BFP-CXCL10$^+$ cells was reconstructed, rendered and registered as a point in space using Imaris and Python. A point (cell centroid) was designated a given radius, R (μm). Each point was defined as a core point within the cluster if it had at least a specified number of neighbor points (N) within its specified radius (R). A point was defined a boarder point if it had fewer than N points within R, but had at least one core point in its radius. A noise point was any point that was not a core point nor a border point, and therefore not part of a cluster. **(B)** BFP-CXCL10$^+$ cell cluster (blue) within the d5 OVA/CFA immunized REX3 dermis, by IV-MPM: gray, SHG; scale bar, 50μm. In processed image, BFP-CXCL10$^+$ cells were reconstructed and volumetrically rendered along with a cluster surface rendering in 3D using Imaris. **(C)** Combinations of different R and N were plotted using a grid-search approach to find inflection points after which return on metric of choice is incremental. **(D)** Visual examples of different N cutoffs with R=50 μm, using DBSCAN-based algorithm in Python. **(E)** BFP-CXCL10$^+$ cell clusters (blue) within d5 OVA/CFA immunized REX3 dermis, tiled image composed of 18 imaging fields of 500 (x) x 500 (y) x 60 (z) μm by IV-MPM: CD31$^+$ vessels, red; tiled image scale bar, 200μm. **(F)** DBSCAN-based algorithm was used for 3D cluster identification (multi-color) and reconstruction, using Python; R=50 μm, N=10. **(G)** BFP-CXCL10$^+$ cells (blue) within a naïve REX3 dermis, tiled image composed of 18 imaging fields as in E. **(H)** DBSCAN-based algorithm was used for 3D cluster identification and reconstruction, using Python; R=50 μm, N=10.

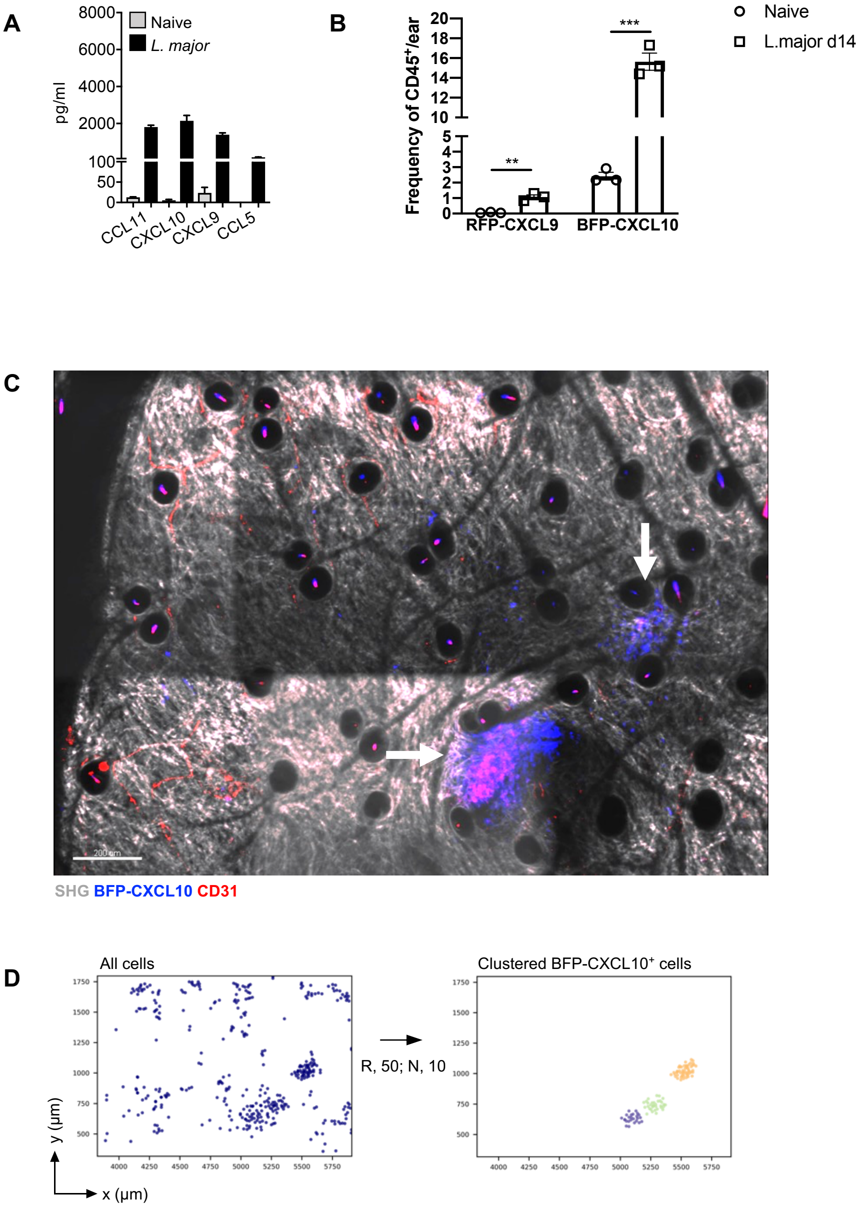


**Figure S3. CXCL10⁺ cell clustering in the *L. major* infected skin**
**(A)** Chemokine protein levels from naïve and 14-day *L. major* infected WT ears, by Luminex. **(B)** frequency of BFP-CXCL10 and RFP-CXCL9 expressing cells within live singlets CD45⁺ cells from 14 day *L. major* infected REX3 ears, by flow cytometry. **(C)** Perivascular clustering of BFP-CXCL10⁺ cells (blue) within 14-day *L. major* infected REX3 dermis, by IV-MPM: scale bar, 200 µm; gray, SHG; anti-CD31 (red) antibody injected i.v. 15 minutes before imaging. **(D)** Cluster analysis using DBSCAN-based algorithm.

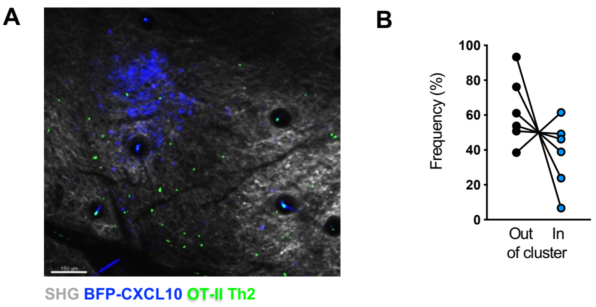


**Figure S4. Th2 cells localization relative to CXCL10⁺ clusters**
**(A)** Adoptively transferred fluorescently labeled OT-II Th2 cell (green) localization to BFP-CXCL10⁺ clusters (blue) within d5 OVA/CFA immunized REX3 ear. Maximal z-projection image by IV-MPM: scale bar, 50μm. **(B)** Percentage of OT-II Th2 cells inside or outside of BFP-CXCL10⁺ clusters in d5 OVA/CFA immunized REX3 dermis, by IV-MPM and Imaris.

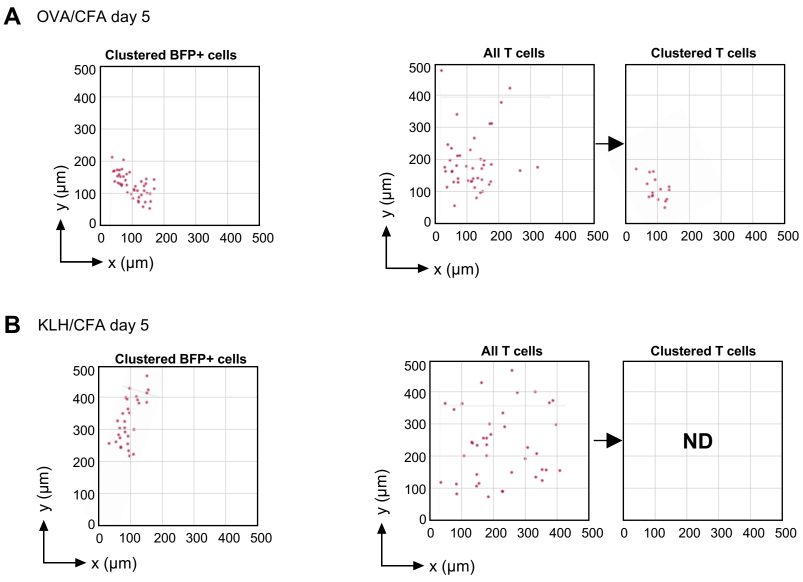


**Figure S5. Antigen-dependent Th1 accumulation in BFP-CXCL10$^+$ clusters**

Adoptively transferred fluorescently labeled OT-II WT Th1 cell localization to BFP-CXCL10$^+$ clusters within d5 OVA/CFA and KLH/CFA immunized REX3 ear. **(A, B)** DBSCAN-based algorithm for semi-automated 3D cluster reconstruction of BFP-CXCL10$^+$ (left) and Th1 cells (right) from OVA/CFA **(A)** and KLH/CFA **(B)** immunized REX3 ears. ND, none detected. Representative analysis of images shown in Fig. 2D.

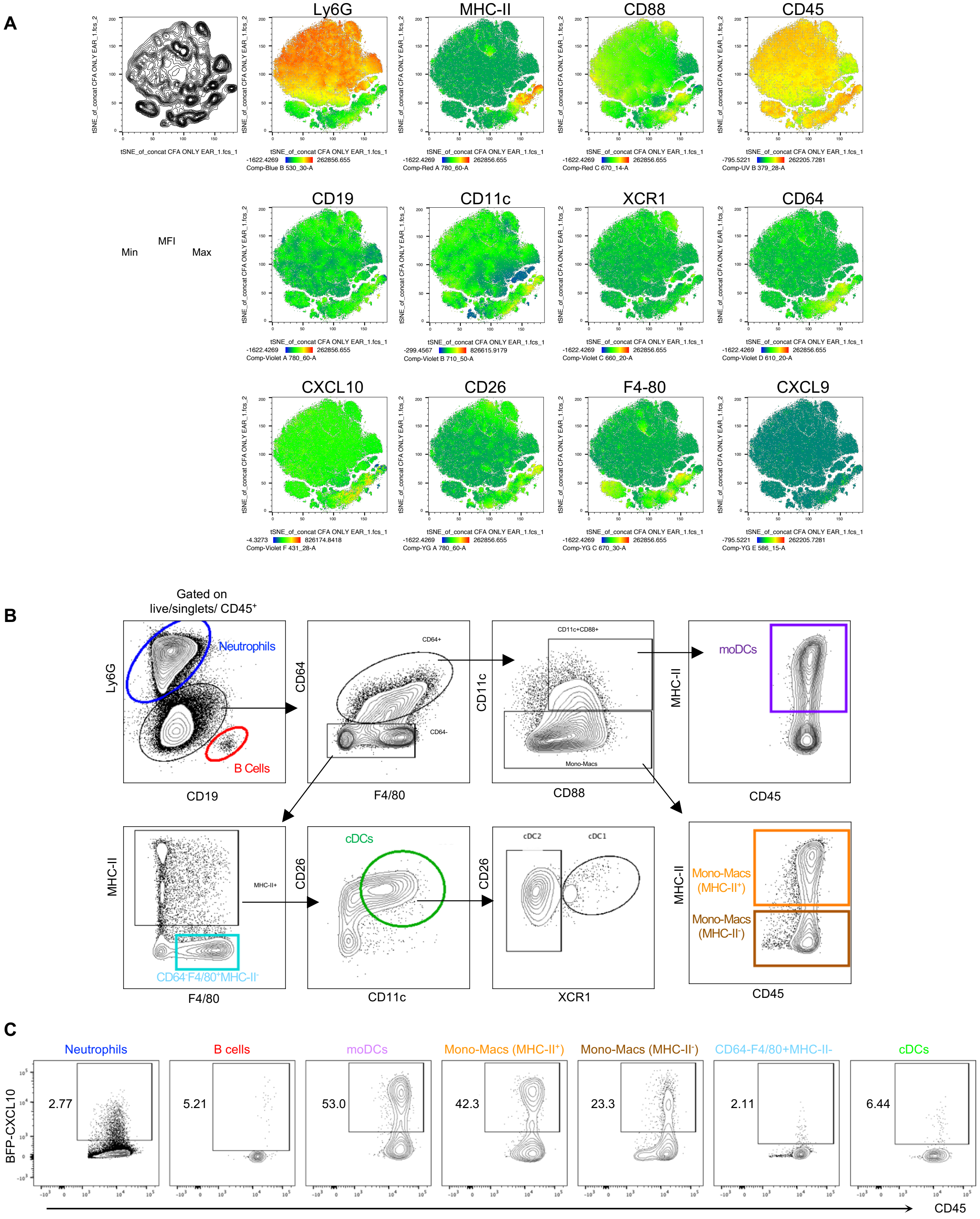


**Figure S6. Gating strategies to define the immune subsets with the inflamed skin**
**(A)** tSNE analysis was performed on singlet live CD45$^{+}$ cells using concatenated multi-color flow cytometry data from d5 OVA/CFA immunized REX3 ears. **(B)** Gating strategy to define the immune subsets within the immunized skin, pre-gated on singlet live CD45$^{+}$ cells: Neutrophils, Ly6G$^{high}$; B cells, CD19$^{+}$; moDCs, Ly6G$^{low}$CD19$^{-}$CD64$^{+}$F4/80$^{+}$CD11c$^{+}$MHC-II$^{+}$; Mono-Macs (MHC-II$^{+}$), Ly6G$^{low}$CD19$^{-}$CD64$^{+}$F4/80$^{+}$CD11c$^{-}$MHC-II$^{+}$; Mono-Macs (MHC-II$^{-}$), Ly6G$^{low}$CD19$^{-}$CD64$^{+}$F4/80$^{+}$CD11c$^{-}$MHC-II$^{-}$; CD64$^{-}$F4/80$^{+}$ MHC-II$^{-}$, Ly6G$^{low}$CD19$^{-}$CD64$^{-}$F4/80$^{+}$MHC-II$^{-}$; cDCs, Ly6G$^{low}$CD19$^{-}$CD64$^{-}$F4/80$^{+/-}$MHC-II$^{+}$CD11c$^{+}$CD26$^{+}$; cDC1, CD26$^{+}$XCR1$^{+}$; cDC2, CD26$^{+}$XCR1$^{-}$. **(C)** Representative plots of BFP-CXCL10 expressing cells within each subset in (B), d5 OVA/CFA immunized REX3 ears.

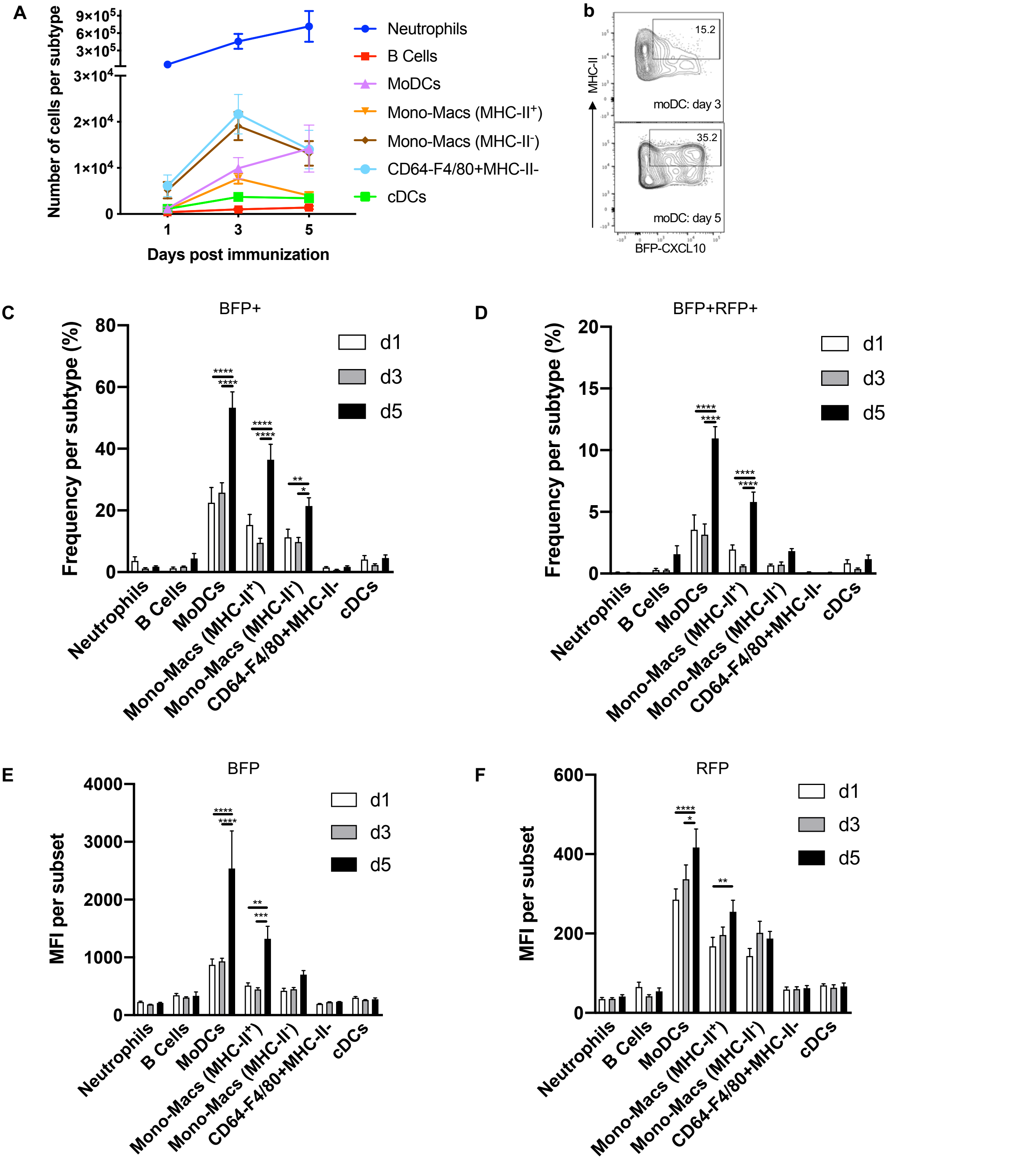


**Figure S7. Kinetic analysis of immune subsets and CXCR3 chemokine expression within the immunized skin**. **(A)** Number of cells per immune subset within live singlets CD45+ cells from OVA/CFA immunized REX3 ears over time. **(B)** BFP-CXCL10 and MHC-II expression within the moDC population (Ly6GlowCD64+F4/80+Cd11c+MHC-II+) in the ear skin of day 3 and day 5 OVA/CFA immunized REX3 mice. **(C)** Frequency of BFP-CXCL10+ cells and **(D)** BFP-CXCL10+RFP-CXCL9+ cells within each subtype from (B). **(E)** BFP-CXCL10 and **(F)** RFP-CXCL9 MFI within each immune subset from (B). Bars represent mean + SEM. Statistics by two-way ANOVA, *p≤0.05, **p≤0.01, ***p≤0.001, ****p≤0.0001.
